# PBRM1-VHL cooperation rewires lipid and iron metabolism to promote ferroptosis resistance in clear cell renal cell carcinoma

**DOI:** 10.64898/2026.05.02.722429

**Authors:** Guanming Jiao, Luisa F Baracaldo-Lancheros, Alisha Dhiman, Christina R Ferreira, Emily C Dykhuizen

## Abstract

Clear cell renal cell carcinoma (ccRCC) is initiated by biallelic loss of the tumor suppressor *VHL* followed by additional genomic alterations, including loss of tumor suppressors PBRM1, BAP1 or SETD2. Although ccRCC is known to be intrinsically sensitive to ferroptosis, the contribution of PBRM1 to this vulnerability, and how it interfaces with VHL loss, has remained unexplored.

Using isogenic ccRCC and non-transformed cell models, we show that re-expression of PBRM1 and/or VHL attenuates GPX4-inhibitor-induced ferroptosis, and that the two tumor suppressors act cooperatively: dual reconstitution confers the greatest protection, suppresses lipid peroxidation, and preserves redox homeostasis. Time-resolved RNA-seq reveals that PBRM1 and VHL establish additive, largely non-overlapping "ground-state" transcriptional programs. Integrated pharmacogenomic, CRISPR dependency, and lipidomic analyses converge on two protective axes: restricted labile iron and MUFA-biased membrane remodeling. These findings identify PBRM1 as a previously unrecognized modulator of ferroptosis and define a cooperative chromatin–metabolic axis that buffers ferroptotic cell death in ccRCC.

## Introduction

Ferroptosis is a regulated form of cell death that has recently emerged as a key vulnerability in cancer cells with altered metabolism and redox balance (1). Unlike other types of regulated cell death (RCD), ferroptosis is governed by the interplay between phospholipid-containing polyunsaturated fatty acids (PUFA) susceptible to peroxidation, antioxidant systems such as glutathione peroxidase 4 (GPX4), and cellular iron availability (Fig. 1). Disruption of this balance by inhibiting GPX4 or altering lipid and iron metabolism triggers catastrophic membrane damage and cell death (2). Because ferroptotic sensitivity is tightly linked to cellular metabolic state, ferroptosis represents an attractive vulnerability in metabolically rewired tumors, particularly those exhibiting altered lipid metabolism. PUFA incorporation into membrane phospholipids creates substrates for lipid peroxidation, whereas enrichment of monounsaturated fatty acids (MUFAs) and neutral lipid storage can buffer membranes against oxidative damage by limiting PUFA availability (3). Iron metabolism provides an additional layer of regulation: labile iron pools fuel lipid peroxidation through Fenton chemistry, whereas iron sequestration and trafficking pathways modulate the threshold for ferroptotic death (4).

**Figure 1.**
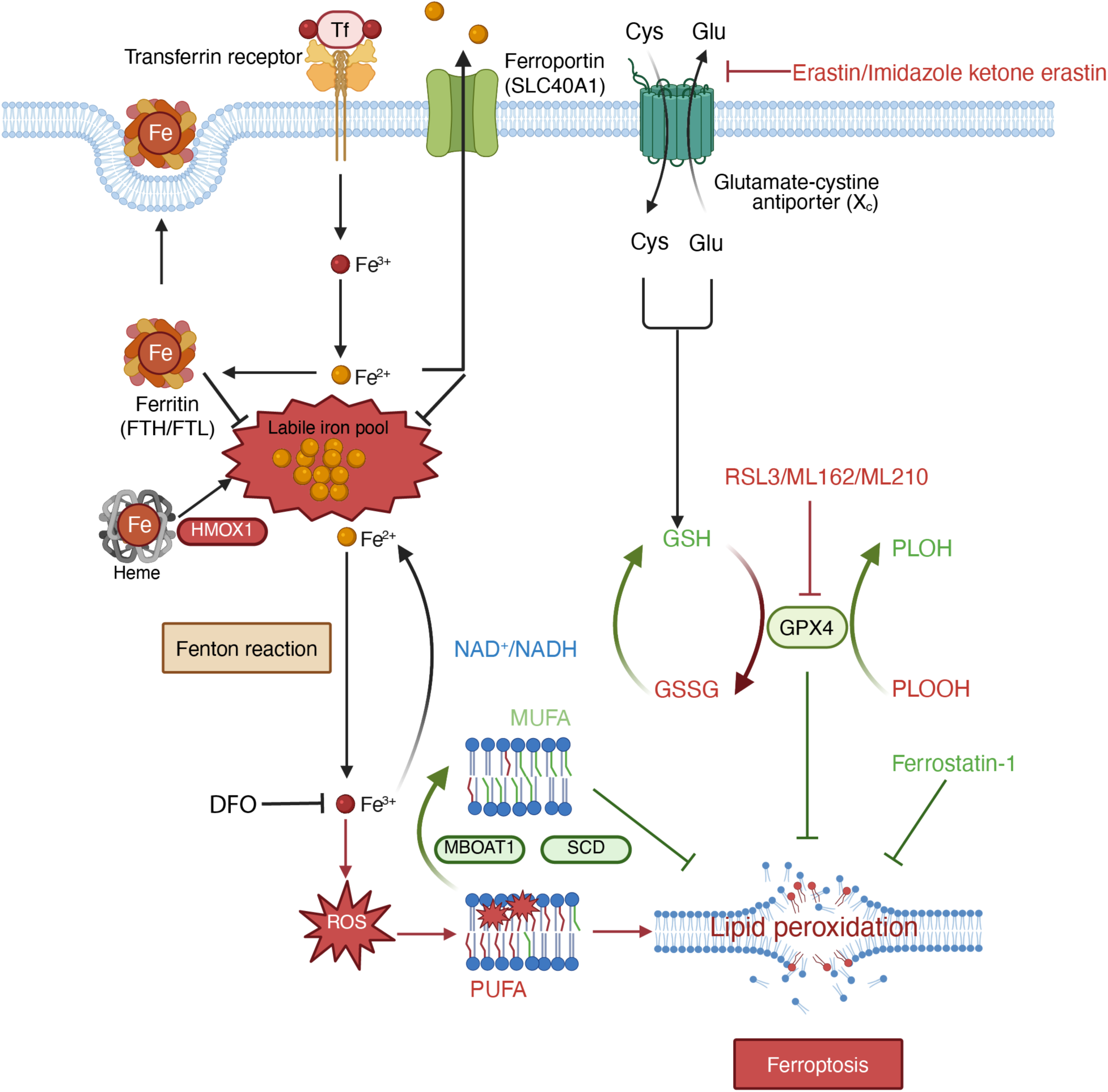
Overview of the ferroptosis regulatory network. Ferroptosis arises from three interacting axes: iron handling (iron uptake via transferrin receptor, storage in ferritin (FTH1/FTL), export via ferroportin/SLC40A1, and heme catabolism by HMOX1), antioxidant defense (system Xc⁻-dependent cystine import supporting GSH synthesis and GPX4-mediated reduction of phospholipid hydroperoxides (PLOOH) to PLOH), and membrane lipid composition (PUFA substrates versus MUFA-enriched, peroxidation-resistant phospholipids generated by SCD and MBOAT1). Fenton-chemistry-driven Fe²⁺ ROS oxidizes PUFA-containing phospholipids, leading to ferroptotic death. Pharmacological perturbations are indicated: erastin/IKE (system Xc⁻), RSL3/ML162/ML210 (GPX4), Deferoxamine (DFO, iron chelation) and ferrostatin-1 (radical trapping).

Clear cell renal cell carcinoma (ccRCC), the most common subtype of kidney cancer, is characterized by alterations in lipid and iron metabolism (5). Hallmark features of ccRCC include extensive lipid droplet accumulation, dysregulated fatty acid handling, and altered oxidative metabolism, largely driven by loss of the von Hippel-Lindau (VHL) tumor suppressor and consequent activation of hypoxia-inducible factor (HIF) signaling (6). Recent studies have shown that HIF signaling enriches PUFA content, and that the restoration of VHL can reprogram metabolism and suppress lipid peroxidation (7,8). As a result, ccRCC cell lines exhibit increased sensitivity to ferroptosis-inducing agents (7,9). Although ferroptosis is an appealing therapeutic strategy for cRCC, there is considerable variation between tumors, highlighting the need to further understand how these lipid and iron regulatory axes are coordinated, particularly in the context of different genetic backgrounds.

Beyond VHL loss, ccRCC is characterized by frequent inactivation of chromatin-associated genes. *PBRM1*, which encodes a subunit of the PBAF SWI/SNF chromatin remodeling complex, is the second most commonly mutated gene in ccRCC (10), while *SETD2* and *BAP1*, which are involved in histone modification, are the third and fourth most commonly mutated genes (11).

PBRM1 loss occurs in approximately 40% of ccRCC tumors, primarily in conjunction with VHL deletion; in mouse models both *PBRM1* and *VHL* loss are required for tumorigenesis (12,13). PBRM1’s role as a tumor suppressor has been attributed to transcriptional programs controlling cell identity, metabolism, and stress responses (14–17), and to roles in genomic stability and DNA damage repair (18). While VHL, SETD2, and BAP1 have been demonstrated to provide resistance to ferroptosis (7,8,19,20), the role of PBRM1 in ferroptotic susceptibility has not been examined. Moreover, how PBRM1 cooperates with VHL-dependent metabolic programs to influence lipid and iron homeostasis in ccRCC has not been investigated.

Here, we investigate how PBRM1 and VHL individually and cooperatively regulate ferroptosis sensitivity in ccRCC. Using genetically defined cell models, we demonstrate that restoration of both tumor suppressors confers maximal resistance to GPX4 inhibitor-induced ferroptosis, exceeding the protective effects observed with either protein alone. Through integrated transcriptomic and lipidomic analyses, we show that PBRM1-VHL cooperation rewires lipid and iron metabolic pathways, establishing a metabolic state characterized by reduced lipid peroxidation stress and altered iron handling. These findings identify PBRM1 as a previously unrecognized modulator of ferroptosis and reveal a cooperative tumor suppressor network that links chromatin regulation to metabolic control of cell death in ccRCC.

## Results

### Clear-cell renal carcinoma exhibits intrinsic GPX4 dependency and ferroptosis sensitivity

GPX4 is a key antioxidant enzyme that suppresses lipid peroxidation and protects cells from ferroptosis (21). To investigate the overall sensitivity of kidney cancer cells to ferroptosis, we calculated the mean *GPX4* dependency score for each tumor lineage using the genome-scale CRISPR screening data from DepMap (Chronos 25Q3+) (https://depmap.org/portal/). Kidney cancer cell lines ranked among those with the most negative *GPX4* dependency scores, indicating an increased reliance on the GPX4 pathway for survival (Figs. 2A, B). Consistent with this dependency, pharmacogenomic profiling from the Cancer Therapeutics Response Portal (CTRP) (https://portals.broadinstitute.org/ctrp.v2/) revealed that kidney cancer lineages were among the most sensitive to three structurally distinct GPX4 inhibitors (RSL3, ML162, ML210), displaying the lowest median AUC values compared to non-kidney lineages (Figs. 2C, D).

**Figure 2.**
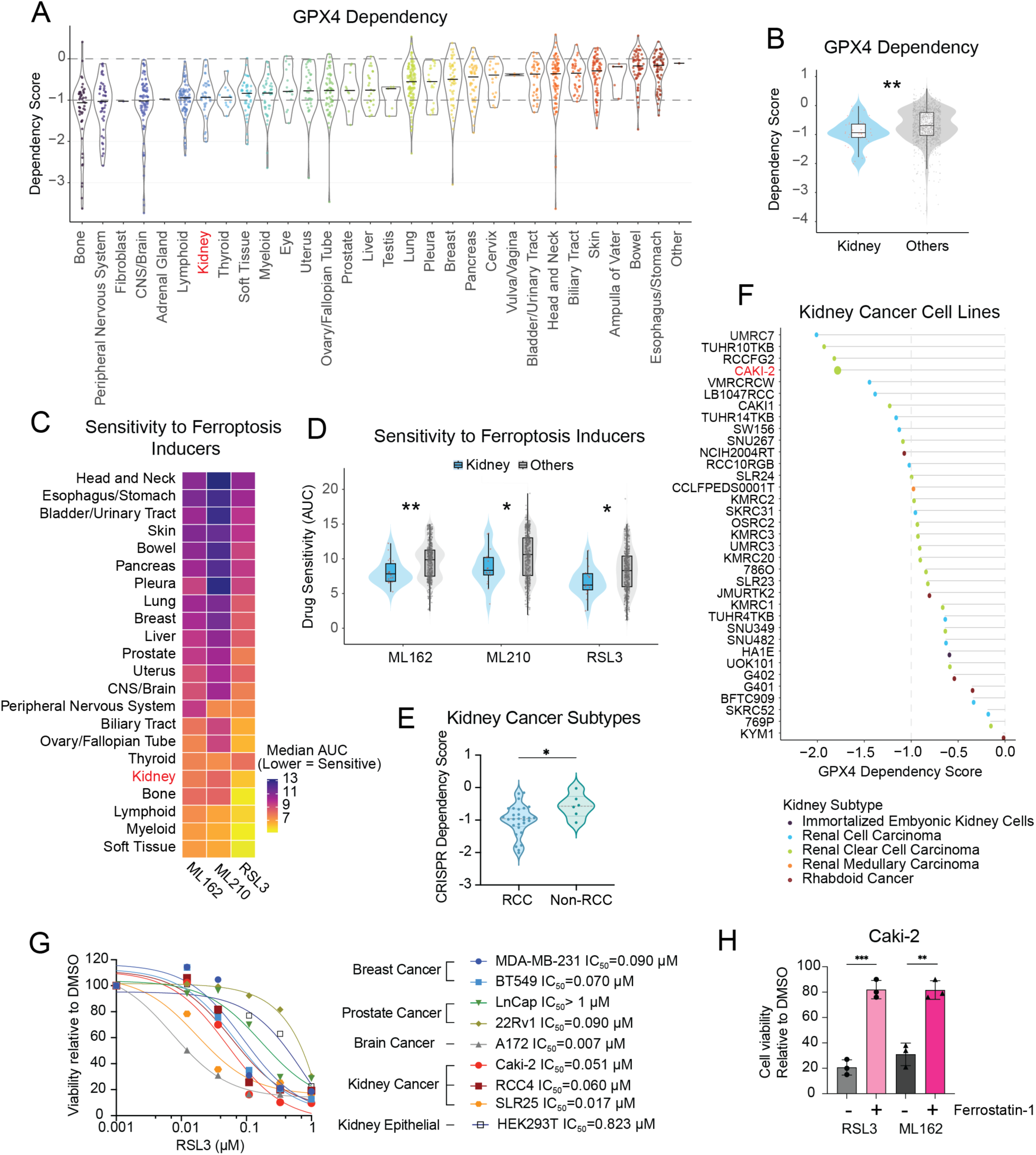
Clear-cell renal carcinoma exhibits intrinsic GPX4 dependency and ferroptosis sensitivity. (A) GPX4 CRISPR dependency (Chronos scores) across tumor lineages from DepMap Public 25Q3+. Individual points represent cell lines; distributions are displayed as violin plots. More negative scores indicate greater dependency on GPX4. Kidney lineage is highlighted in red. (B) Comparison of GPX4 dependency scores for kidney lineage cell lines (*n* = 36) versus all other tumor lineages (*n* = 822). Statistical significance was determined using a two-sided Wilcoxon rank-sum test (\*\**P* ≤ 0.01). (C) Heatmap summarizing median drug response (AUC) across tumor lineages from CTRP for the GPX4 inhibitors ML162, ML210, and RSL3. Lower AUC indicates greater sensitivity. (D) Distribution of CTRP AUC values for ML162, ML210, and RSL3 comparing kidney lineage cell lines (*n* = 18) versus all other tumor lineages (*n* = 822). Statistical significance was determined using a two-sided Wilcoxon rank-sum test (\**P* ≤ 0.05; \*\**P* ≤ 0.01). (E) GPX4 dependency scores within kidney-derived lines, comparing renal cell carcinoma (*n* = 29) versus non-RCC kidney cell lines (*n* = 5). Statistical significance was determined using a two-sided Wilcoxon rank-sum test (\**P* ≤ 0.05). (F) Kidney cancer cell lines ranked by GPX4 CRISPR dependency score, color-coded by kidney subtype (immortalized embryonic kidney, renal cell carcinoma, renal clear cell carcinoma, renal medullary carcinoma, rhabdoid cancer). Caki-2 is highlighted in red. Dashed line indicates a dependency score of −1. (G) Dose-response viability curves following RSL3 treatment (16 h) in indicated cell lines from breast (MDA-MB-231, BT549), prostate (LNCaP, 22Rv1), brain (A172), kidney cancer (Caki-2, RCC4, SLR25), and normal kidney epithelial (HEK293T) origins. Viability is normalized to DMSO-treated controls. IC₅₀ values were derived from 4-parameter nonlinear regression on *n* = 3 biological replicates. IC₅₀ values are indicated. (H) Ferrostatin-1 (Fer-1; 1 µM) rescues RSL3 (0.33 µM)- and ML162 (0.33 µM)-induced cytotoxicity in Caki-2 cells following 18 h treatment. Viability is normalized to DMSO-treated controls. (G-H) Data are presented as mean ± SD (*n* = 3 biological replicates). Statistical significance was assessed by unpaired two-sided Student’s t-test (\**P* ≤ 0.05; \*\**P* ≤ 0.01; \*\*\**P* ≤ 0.001).

Subtype annotation revealed that renal cell carcinoma (RCC) lines exhibited stronger GPX4 dependencies than non-RCC kidney lines (Fig. 2E), consistent with prior reports identifying clear-cell carcinomas as ferroptosis-sensitive (8). Ranking kidney cancer cell lines by GPX4 dependency further enriched for clear-cell RCC models, with Caki-2 among the strongest dependencies (Fig. 2F).

We experimentally validated this vulnerability using representative ccRCC models (Caki-2, RCC4, SLR25) treated with the GPX4 inhibitor RSL3 and compared responses to non-RCC cancer cell lines. RCC models displayed significantly lower IC₅₀ values than most non-RCC lines (Fig. 2G), except for the glioblastoma line A172, consistent with known CNS GPX4 dependency (Fig. 2A). Sensitivity was confirmed using a second GPX4 inhibitor, ML162, and ferrostatin-1 (Fer-1), a lipophilic radical-trapping antioxidant that blocks phospholipid peroxidation, fully rescued RSL3- and ML162-induced cytotoxicity (Fig. 2H), confirming ferroptotic cell death. Together, these analyses identify ccRCC as a GPX4-dependent lineage with intrinsic ferroptosis sensitivity.

### PBRM1 and VHL cooperatively regulate GPX4 dependency and ferroptosis sensitivity in ccRCC

Given the intrinsic vulnerability of ccRCC to GPX4 inhibition, we next examined how VHL and PBRM1 modulate ferroptosis sensitivity. Using integrated DepMap genomic annotations, renal carcinoma cell lines were stratified by VHL and PBRM1 status. VHL and VHL/PBRM1 loss corresponded to the strongest GPX4 dependencies, whereas PBRM1 loss alone had a more modest effect (Fig. 3A) with effect-size analysis identifying VHL loss as the dominant determinant (Fig. 3B).

**Figure 3.**
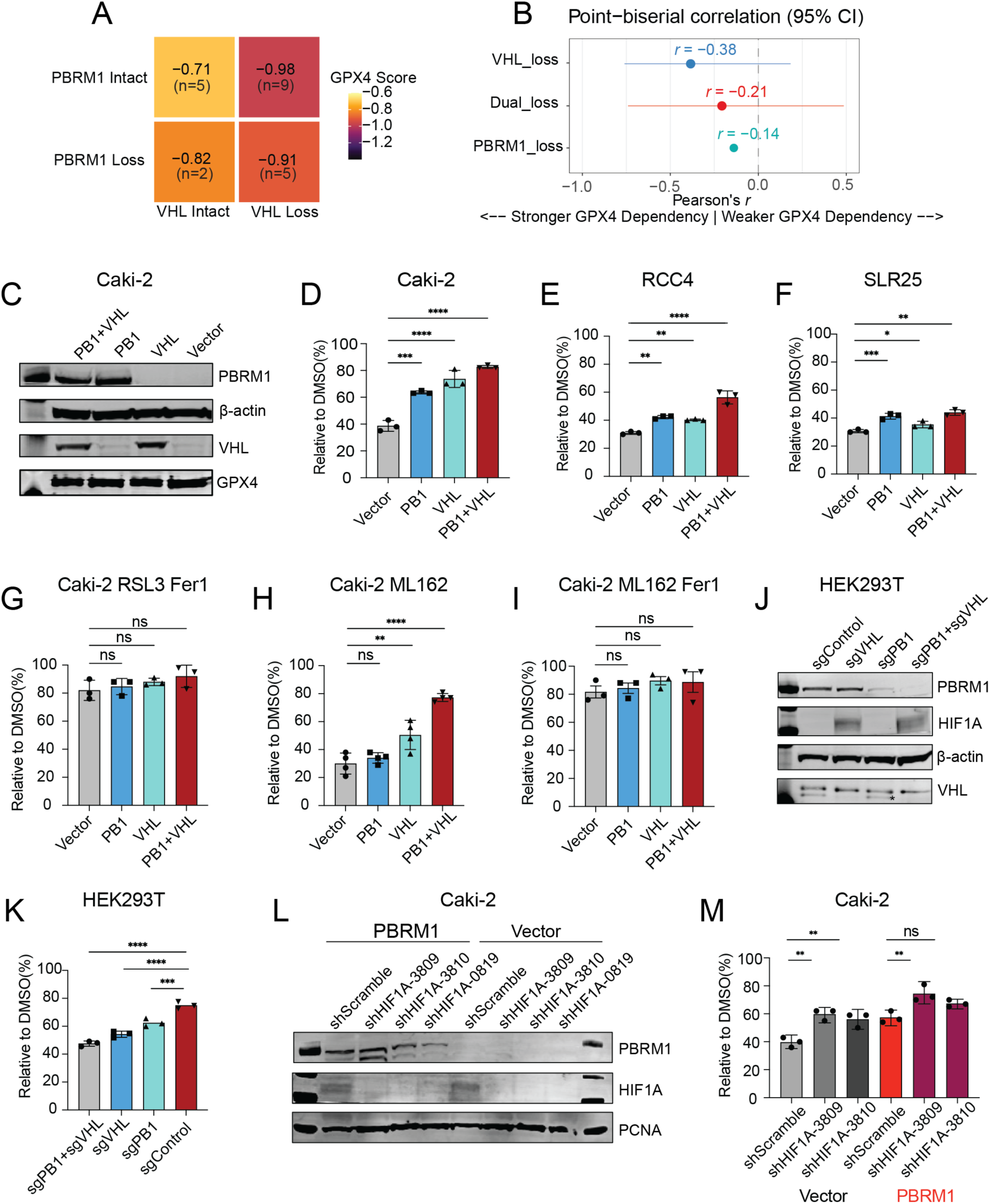
PBRM1 and VHL cooperatively regulate GPX4 dependency and ferroptosis sensitivity in ccRCC. (A) Mean GPX4 Chronos dependency scores (from DepMap) for renal carcinoma cell lines stratified by PBRM1 and VHL status. Values within each cell are the mean dependency score; *n* indicates the number of cell lines per genotype group. (B) Effect sizes for the association between genotype class and GPX4 dependency, summarized as point-biserial Pearson’s *r* with 95% confidence intervals from bootstrap resampling. (C) Immunoblot of PBRM1, VHL, and GPX4 in Caki-2 cells reconstituted with vector, PBRM1, VHL, or PBRM1+VHL. β-actin serves as a loading control. (D-F) Viability of Caki-2 (D), RCC4 (E), and SLR25 (F) reconstituted cells following 18 h treatment with RSL3 (0.33 µM for Caki-2 and SLR25; 0.11 µM for RCC4). Viability is normalized to DMSO-treated controls within each genotype. (G) Ferrostatin-1 (Fer-1; 1 µM) rescue of RSL3 (0.33 µM, 18 h)-induced cytotoxicity in Caki-2 reconstituted cells; viability is normalized to DMSO-treated controls within each genotype. (H) Viability of Caki-2 reconstituted cells following treatment with the independent GPX4 inhibitor ML162 (0.33 µM, 18 h), normalized to DMSO-treated controls. (I) Fer-1 (1 µM) rescue of ML162 (0.33 µM, 18 h)-induced cytotoxicity in Caki-2 reconstituted cells; viability is normalized to DMSO-treated controls within each genotype. (J) Immunoblot validation of CRISPR-mediated disruption of PBRM1 and/or VHL in HEK293T cells (sgControl, sgVHL, sgPBRM1, sgPBRM1+sgVHL). HIF1A accumulation in sgVHL and sgPBRM1+sgVHL lanes serves as a functional readout of VHL disruption. β-actin serves as a loading control; asterisk indicates the VHL band. (K) Viability of HEK293T CRISPR-edited cells following RSL3 treatment (0.33 µM, 18 h), normalized to DMSO-treated controls. (L) Immunoblot validation of HIF1A knockdown by independent shRNAs in Caki-2 vector and PBRM1-reconstituted cells; PCNA serves as a loading control. (M) Viability following GPX4 inhibition (RSL3; 0.33 µM, 18 h) in Caki-2 vector and PBRM1-reconstituted cells with HIF1A knockdown, normalized to DMSO-treated controls. (D-I, K, M) Data are presented as mean ± SD with individual replicates shown (*n* = 3 biological replicates). Statistical significance was assessed by one-way ANOVA with Dunnett’s multiple-comparisons test using the indicated reference group (vector for D-I, K; shScramble within each genotype for M). \**P* ≤ 0.05; \*\**P* ≤ 0.01; \*\*\**P* ≤ 0.001; \*\*\*\**P* ≤ 0.0001.

To test causality, we generated isogenic ccRCC cell models (Caki-2, RCC4, SLR25) in which PBRM1, VHL, or both were re-expressed in VHL/PBRM1-deficient backgrounds.

Immunoblotting confirmed protein restoration (Figs. 3C, S1A, B), and GPX4 protein levels were comparable across genotypes (Figs. 3C, S1B). Re-expression of either tumor suppressor significantly attenuated RSL3-induced cytotoxicity, with dual PBRM1+VHL restoration consistently conferring the strongest protection (Figs. 3D–F). Fer-1 fully rescued cell viability and eliminated genotype-dependent differences (Fig. 3G), confirming ferroptosis as the mode of death. The phenotype was confirmed with an independent GPX4 inhibitor, ML162 (Fig. 3H, I).

To assess generality, we disrupted PBRM1 and/or VHL in non-cancerous HEK293T cells using CRISPR-Cas9 (Fig. 3J). Loss of either gene increased sensitivity to GPX4 inhibition, with dual loss producing the strongest effect (Fig. 3K). Similarly, PBRM1 depletion in BT549 breast cancer cells increased RSL3 sensitivity (S1C), indicating a broader role for PBRM1 in ferroptosis resistance.

Finally, we tested whether PBRM1-dependent ferroptosis resistance requires HIF1A (Hypoxia Inducible Factor 1 Subunit Alpha). While HIF1A depletion phenocopied the protective effect of VHL restoration, it did not alter PBRM1-dependent resistance (Figs. 3L, M), indicating that PBRM1 acts independently of HIF1A. These results demonstrate that VHL and PBRM1 cooperatively suppress ferroptosis through additive HIF-dependent and HIF-independent mechanisms.

### PBRM1 and VHL suppress lipid peroxidation and preserve redox homeostasis during ferroptotic stress

To determine whether genotype-dependent protection from GPX4 inhibition reflects biochemical suppression of ferroptosis, we measured lipid peroxidation, the defining executioner signal of ferroptosis. Using the oxidation-sensitive PUFA analog probe C11-BODIPY581/591 in Caki-2 cells, RSL3 treatment induced a robust increase in oxidized fluorescence relative to DMSO controls (Fig. 4A). Re-expression of PBRM1 or VHL significantly attenuated this response, while dual reconstitution provided the strongest suppression, approaching baseline levels (Fig. 4A).

**Figure 4.**
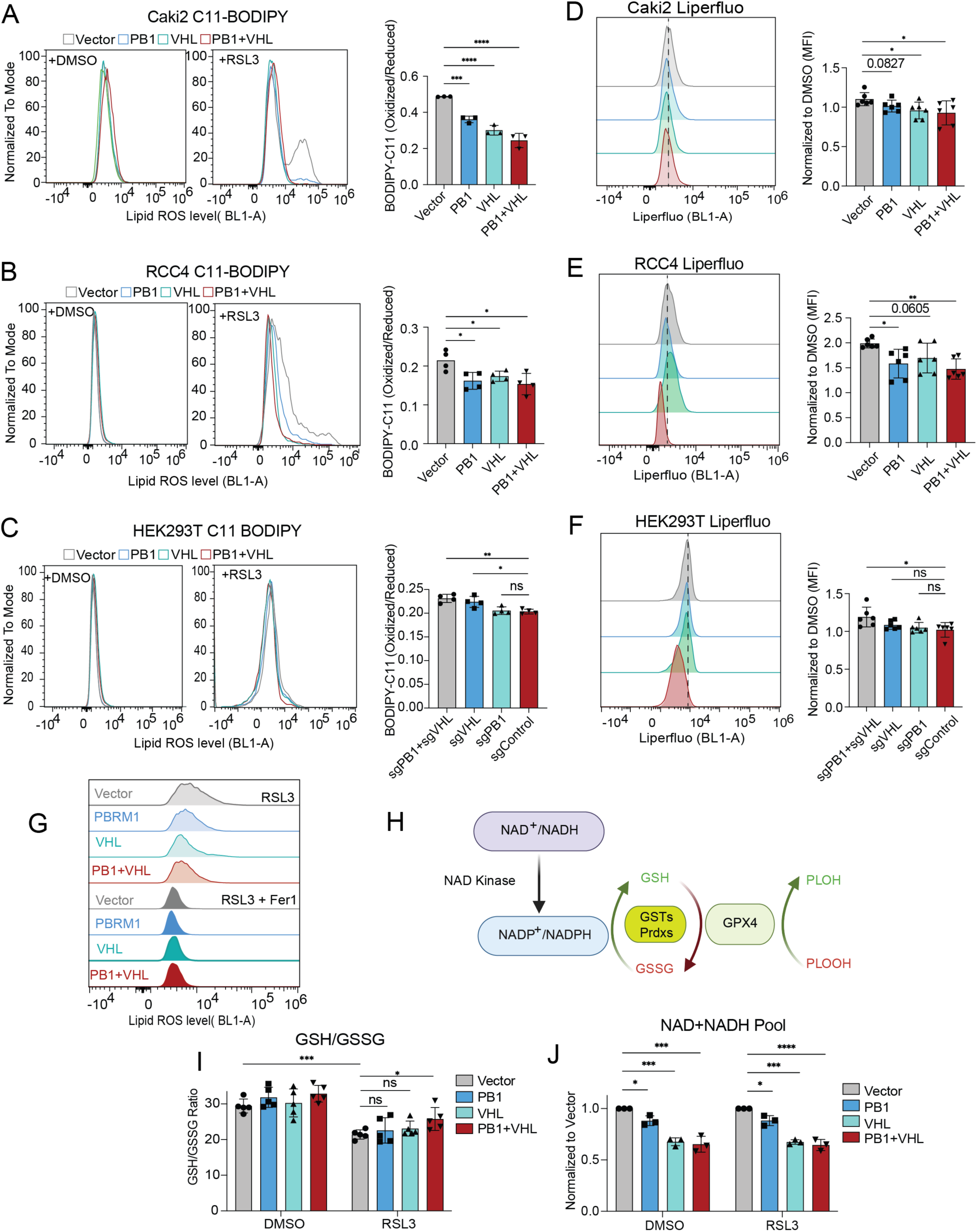
PBRM1 and VHL suppress lipid peroxidation and preserve redox homeostasis during ferroptotic stress. (A) Representative flow cytometry histograms and quantification of BODIPY 581/591 C11 oxidation ratio (oxidized/reduced) in Caki-2 cells expressing vector, PBRM1, VHL, or PBRM1+VHL treated with DMSO or RSL3 (0.33 µM) for 12 hours. (B) As in (A), BODIPY 581/591 C11 oxidation ratio in RCC4 cells reconstituted cells treated DMSO or RSL3 (0.11 µM) for 12 hours. (C) BODIPY 581/591 C11 oxidation ratio in CRISPR-edited HEK293T cells (sgControl, sgPBRM1, sgVHL, sgPBRM1+sgVHL) treated with DMSO or RSL3 (0.33 µM) for 8 hours. (D-E) Representative histograms and mean fluorescence intensity (MFI) quantification of Liperfluo staining in Caki-2 (D) and RCC4 (E) cells following RSL3 treatment as in (A-B). (F) Liperfluo fluorescence as mean fluorescence intensity in CRISPR-edited HEK293T cells following RSL3 treatment as in (C). (G) Representative flow cytometry histograms of BODIPY 581/591 C11 oxidation in Caki-2 reconstituted cells treated with RSL3 (0.33 µM) ± ferrostatin-1 (Fer-1; 1 µM) for 8 h. (H) Schematic illustrating the relationship between the NAD⁺/NADH pool, NADP⁺/NADPH generation via NAD kinase, GSH/GSSG redox cycling, and GPX4-dependent detoxification of lipid peroxides. GSTs: Glutathione S-transferases; Prdxs: Peroxiredoxins. (I) Glutathione redox status (GSH/GSSG ratio) in Caki-2 cells treated with DMSO or RSL3 (0.33 µM) for 8 hours, quantified using the GSH/GSSG-Glo™ Assay. (J) Total NAD (NAD⁺ + NADH) in Caki-2 cells treated with DMSO or RSL3 (0.33 µM) for 8 hours, quantified using the NAD/NADH-Glo™ Assay. Data are presented as mean ± SD with individual replicates shown (*n* = 3 biologically independent replicates for panels A, J, I; *n* = 4 for panels B, C; *n* = 5 for panels I; *n* = 6 for panels (D, E, F). For panels comparing ≥3 genotype groups, statistical significance was assessed by one-way ANOVA with Dunnett’s multiple-comparisons test using vector as the reference group. For panels I and J, comparisons between DMSO and RSL3 within each genotype were assessed by unpaired two-sided Student’s *t*-test. Exact *P*-values are shown in the figure for comparisons that did not reach the threshold for asterisk annotation. \**P* ≤ 0.05; \*\**P* ≤ 0.01; \*\*\**P* ≤ 0.001; \*\*\*\**P* ≤ 0.0001.

This pattern was reproduced in RCC4 cells (Fig. 4B) and in HEK293T cells following CRISPR-mediated loss of PBRM1 and/or VHL (Fig. 4C). Lipid peroxide suppression was confirmed using the oxidation-sensitive orthogonal probe Liperfluo (Figs. 4D–F). Co-treatment with ferrostatin-1 (Fer-1) eliminated genotype-dependent differences in C11-BODIPY oxidation (Fig. 4G), confirming that differences reflected ferroptotic lipid peroxidation rather than nonspecific oxidative stress.

Because GPX4 detoxifies lipid peroxides by consuming cellular reducing equivalents (Fig. 4H), we next assessed redox buffering during GPX4 inhibition. In untreated cells, PBRM1/VHL re-expression does not alter the ratio of GSH to GSSG. Following extended RSL3 treatment, vector control cells exhibited a reduced GSH/GSSG ratio, whereas PBRM1 and VHL-reconstituted cells retained a slightly higher ratio (Fig. 4I), consistent with diminished lipid peroxide burden. In parallel, PBRM1- and VHL-expressing cells displayed reduced total NAD⁺+NADH pools under both basal and RSL3-treated conditions (Fig. 4J), indicating lower oxidative demand. Together, these results show that PBRM1 and VHL limit lipid peroxidation and oxidative stress, reducing reliance on GPX4 for redox homeostasis.

### PBRM1 and VHL establish transcriptional states that constrain the RSL3 response

Because PBRM1 regulates transcription through chromatin remodeling, and VHL regulates transcription through HIFα degradation, we examined whether ferroptosis resistance arises from altered basal transcription or from differential responses to GPX4 inhibition. Caki-2 cells were profiled following RSL3 treatment at an early (4 h) and a later (10 h) time point. At 4 h, only vector control cells mounted a significant transcriptional response (66 DEGs, Padj < 0.05), whereas PBRM1-, VHL-, and PBRM1+VHL-reconstituted cells showed minimal changes (Fig. 5A). By 10 h, all genotypes exhibited similar transcriptional responses whose magnitude correlated with ferroptosis sensitivity (Fig. 5B). RSL3-induced gene expression changes were directionally conserved across genotypes (Pearson *r* = 0.48-0.55), but the magnitude of induction was attenuated in PBRM1/VHL reconstituted cells, as reflected by shallower regression slopes compared to one from empty vector control cells (Fig. 5C). Principal component analysis identified genotype as the dominant source of variance, with RSL3 treatment producing secondary effects (Fig. 5D). Consistent with this, baseline transcriptional changes driven by PBRM1 and/or VHL were preserved following GPX4 inhibition (Figs. 5E, S2A, B).

**Figure 5:**
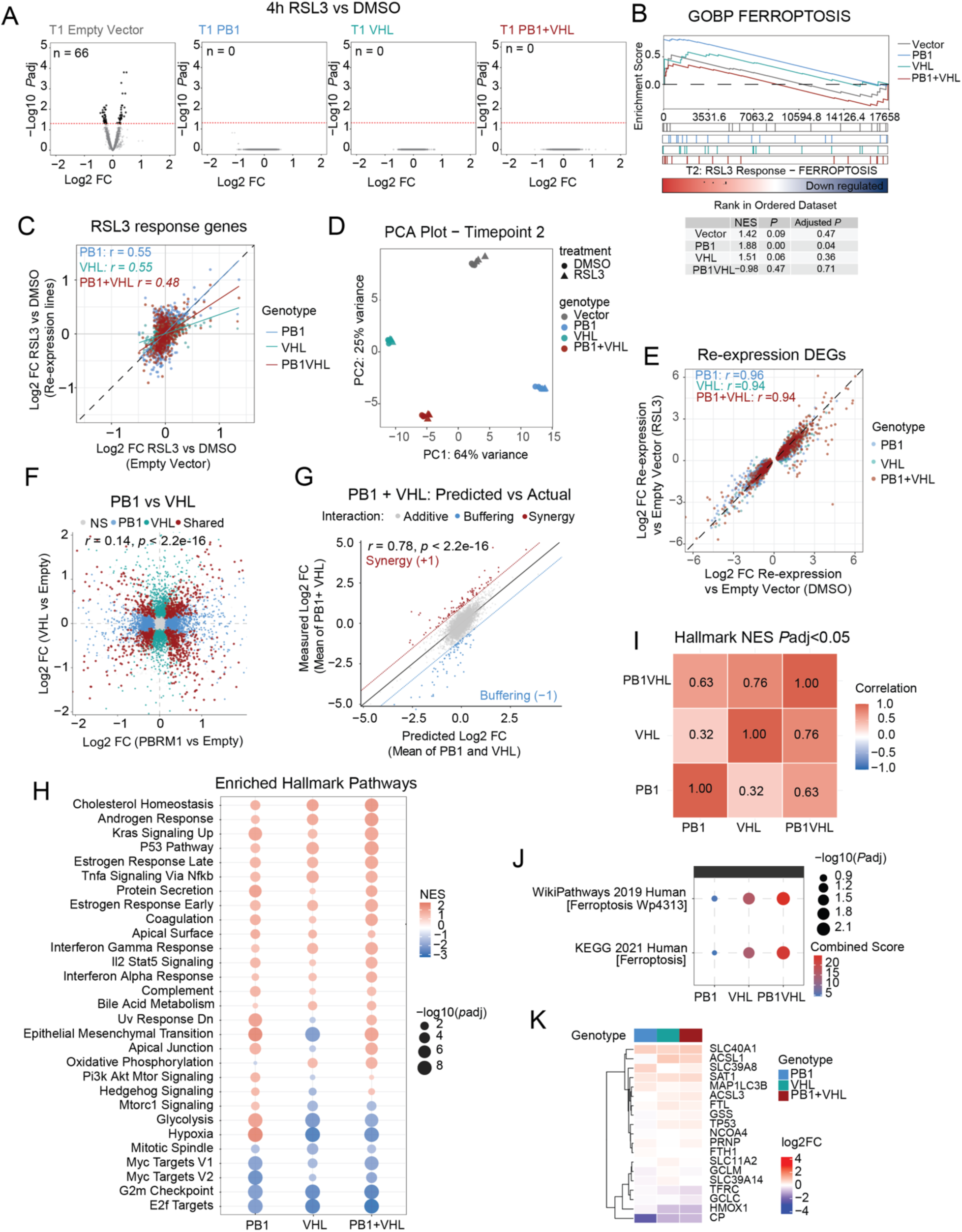
PBRM1 and VHL establish transcriptional states that constrain the RSL3 response. (A) Volcano plots of differentially expressed genes (DEGs) in Caki-2 cells expressing vector (Empty), PBRM1, VHL, or PBRM1+VHL following RSL3 treatment (0.33 μM, 4 h; T1). The number of statistically significant DEGs (adjusted *P* < 0.05) is indicated for each genotype. (B) Gene Set Enrichment Analysis (GSEA) of the GO Biological Process ferroptosis gene set (GO: BP FERROPTOSIS) comparing RSL3 versus DMSO at T2 (10 h) across all four genotypes. Enrichment scores, normalized enrichment scores (NES), nominal *P*-values, and adjusted *P*-values are summarized in the accompanying table. (C) Scatter plots of RSL3-induced log2 fold-changes at T2 in PBRM1-, VHL-, and PBRM1+VHL-reconstituted cells compared to Empty vector cells, with Pearson correlation coefficients (*r*) indicated. Each point represents an individual gene. (D) Principal component analysis (PCA) of T2 transcriptomes across all genotypes and treatment conditions (DMSO and RSL3). Each point represents an individual sample; percentage of variance explained by each principal component is indicated on the respective axis. (E) Scatter plots comparing genotype-versus-Empty log2 fold-changes under DMSO and RSL3 conditions at T2 for PBRM1, VHL, and PBRM1+VHL reconstituted cells, with Pearson correlation coefficients (*r*) indicated. Each point represents an individual gene. (F) Scatter plot comparing PBRM1-versus-Empty and VHL-versus-Empty log2 fold-changes at T2 under DMSO conditions. Points are colored by genotype-specificity classification (PBRM1-specific, VHL-specific, or shared). Global Pearson *r* and shared-gene Pearson *r* are indicated. (G) Epistasis/additive analysis of combined PBRM1 and VHL transcriptional effects at T2 under DMSO conditions. The predicted log2 fold-change for the PBRM1+VHL state was calculated as the mean of individual PBRM1 and VHL log2 fold-changes and plotted against the observed PBRM1+VHL log2 fold-change. Points are colored by interaction class additive, buffering, or synergy (threshold = 1). Pearson *r* = 0.78, *P* < 2.2 × 10⁻¹⁶. (H) Hallmark gene set enrichment analysis (GSEA) bubble plot summarizing pathway-level transcriptional differences for each reconstituted genotype relative to Empty vector under DMSO treatment at T2. Bubble size represents -log10 (adjusted *P*) and color represents NES. Only pathways with adjusted *P* < 0.05 are shown. (I) Pairwise correlation matrix of Hallmark NES profiles between genotypes (PBRM1, VHL, PBRM1+VHL) under DMSO conditions at T2. Color intensity reflects Pearson correlation coefficient. (J) Over-representation analysis (ORA) using EnrichR of ferroptosis-related gene sets from KEGG 2021 Human and WikiPathways 2019 Human databases for each reconstituted genotype under DMSO condition at T2. Bubble size represents -log10(adjusted *P* value) and fill color represents combined enrichment score. (K) Heatmap of log2 fold-changes for leading-edge ferroptosis-associated genes identified in (J), across genotypes of DMSO conditions at T2.

To determine whether PBRM1 and VHL operate through shared or distinct programs, we compared genotype-specific transcriptional effects. Transcriptional outputs driven by PBRM1 and VHL were weakly correlated (Fig. 5F), and the PBRM1+VHL state closely followed an additive model (Fig. 5G), indicating that enhanced resistance arises from superposition of largely orthogonal modules rather than synergy.

Pathway analysis identified ferroptosis-relevant programs including hypoxia, mTORC1 signaling, glycolysis, EMT, and iron metabolism (Figs. 5H, S2C). Pairwise correlation of Hallmark NES profiles further showed that PBRM1 and VHL produced largely distinct pathway signatures (*r* = 0.32), with the dual state more closely resembling VHL than PBRM1 (*r* = 0.76 vs 0.63; Fig. 5I). VHL re-expression predominantly suppressed HIF-associated programs, whereas PBRM1 selectively enriched EMT-associated fibrosis and extracellular matrix gene sets (S2D–F). When focusing on ferroptosis-associated pathways, all re-expression states-particularly VHL-were enriched for genes controlling iron handling, glutathione metabolism, lipid remodeling, and autophagy (Fig. 5J, K). These data support a model in which the response to GPX4 is dependent on a pre-existing, genotype-defined ferroptosis-resistant ground state. The PBRM1+VHL program is largely additive, providing a transcriptional framework for the heightened ferroptosis resistance of the double state.

### PBRM1 and VHL restrict cellular iron availability to limit ferroptotic stress

Given the enrichment of iron-handling genes, we examined iron metabolism as a functional axis of resistance. Pathway analysis revealed genotype-specific enrichment of iron regulatory programs (Fig. 6A). Across re-expression states, genes involved in iron storage and efflux (FTH1, FTL, SLC40A1) were upregulated, whereas iron importers (TFRC, SLC11A2) were repressed (Fig. 6B). FTL and SLC40A1 were among the most consistently upregulated genes and CUT&RUN analysis identified PBRM1 and PBAF occupancy at FTL and SLC40A1 promoters (S3A, B). At the protein level, ferritin heavy chain (FTL) expression was robustly increased across all re-expression genotypes (Fig. 6C) with smaller changes in protein expression of FTH1 and TFRC (S3C, D) Functionally, labile Fe²⁺ levels measured by FerroOrange were highest in vector cells and progressively reduced by PBRM1 and VHL re-expression, with the lowest levels in the dual state (Figs. 6D). Total cellular iron (Fe²⁺ + Fe³⁺) followed a similar pattern (Fig. 6E) and neither pattern changed with RSL3 treatment (S3E, F). Iron supplementation potentiated RSL3-induced ferroptosis, whereas iron chelation with DFO preferentially rescued vector and PBRM1-only cells (Fig. 6F). Thus, PBRM1 and VHL constrain ferroptotic stress by limiting iron availability required for lipid peroxidation.

**Figure 6.**
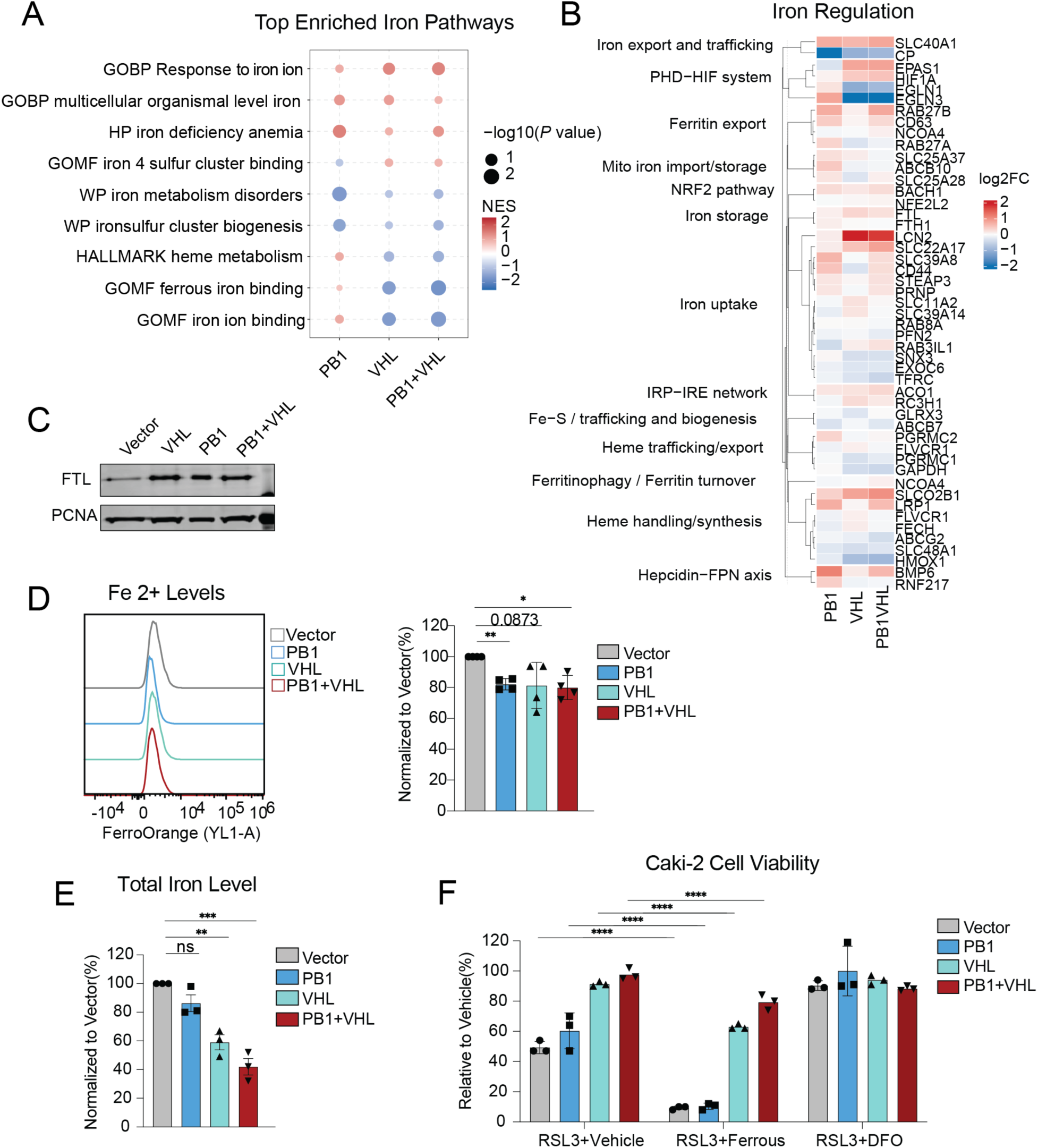
PBRM1 and VHL restrict iron availability through a focused iron-buffering program during GPX4 inhibition. (A) GSEA bubble plot of the top enriched iron-related gene sets across PBRM1, VHL, and PBRM1+VHL reconstituted Caki-2 cells relative to Empty vector under DMSO conditions at T2. Gene sets were drawn from GO Biological Process, GO Molecular Function, Human Phenotype Ontology, WikiPathways, and Hallmark collections. Bubble size represents −log10(*P* value) and fill color represents NES. (B) Heatmap of log2 fold-changes for iron-handling genes across PBRM1, VHL, and PBRM1+VHL reconstituted cells relative to Empty vector under DMSO treatment at T2. Genes are grouped by functional iron-handling category. Color scale represents log2 fold-change. (C) Immunoblot of ferritin light chain (FTL) protein levels in Caki-2 cells expressing Empty vector, VHL, PBRM1, or PBRM1+VHL. PCNA serves as a loading control. (D) Representative flow cytometry histograms (YL1-A channel) and quantification of FerroOrange mean fluorescence intensity (MFI) in Caki-2 cells expressing Empty vector, PBRM1, VHL, or PBRM1+VHL, treated with DMSO or RSL3 (0.33 µM) for 8 h. MFI is normalized to Empty vector within each treatment condition. (E) Total cellular iron content (Fe²⁺ + Fe³⁺) in Caki-2 reconstituted cells under DMSO and RSL3 treatment (0.33 µM, 8 h), quantified by colorimetric iron assay. Values are normalized to Empty vector within each treatment condition. (F) Cell viability in Caki-2 reconstituted cells following RSL3 treatment (0.33 µM, 18 h) in the presence of vehicle (DMSO), ferrous iron supplementation, or the iron chelator deferoxamine (DFO) (100 µM). Viability is expressed relative to vehicle-treated controls within each genotype. Data are presented as mean ± SD with individual replicates shown (*n* = 3 biologically independent replicates). Statistical significance was assessed by one-way ANOVA with Dunnett’s multiple-comparisons test using Empty vector as the reference group. Exact *P*-values are shown in the figure for comparisons that did not reach the threshold for asterisk annotation. \**P* ≤ 0.05; \*\**P* ≤ 0.01; \*\*\**P* ≤ 0.001; \*\*\*\**P* ≤ 0.0001.

### PBRM1 and VHL coordinately shape the lipid metabolic program that supports ferroptosis resistance

Given the central role of lipid composition in ferroptosis, we next analyzed lipid metabolic programs. Enrichment analysis revealed broad regulation of lipid pathways across re-expression states (Fig. 7A). To identify the lipid classes most affected in the VHL/PBRM1 re-expression, we curated 119 genes involved in specific lipid processes (Fig. 7B) (3,22), and found MUFA synthesis and lipophagy to be most highly regulated by PBRM1/VHL. To identify which of these 77 differentially expressed lipid genes are functionally linked to ferroptosis sensitivity, we integrated transcriptional changes with DepMap pharmacogenomic and CRISPR dependency datasets (Fig. 7C). This analysis identified a 13-gene resistance signature spanning MUFA synthesis, phospholipid remodeling, sterol metabolism, redox buffering, and lipid trafficking (Fig. 7D, S4A, B) and a 6-gene sensitivity signature primarily composed of HIF target genes involved in PUFA incorporation (Fig. 7E, S4C, D). Known ferroptosis regulators followed expected patterns using this approach (S4E). To validate functional relevance, pharmacological inhibition of SCD—upregulated across re-expression states—attenuated ferroptosis resistance (S4F), supporting MUFA biosynthesis as a key resistance node. Further, C&R proved the direct occupancy of PBRM1 at these gene promoters (S4G, H).

**Figure 7.**
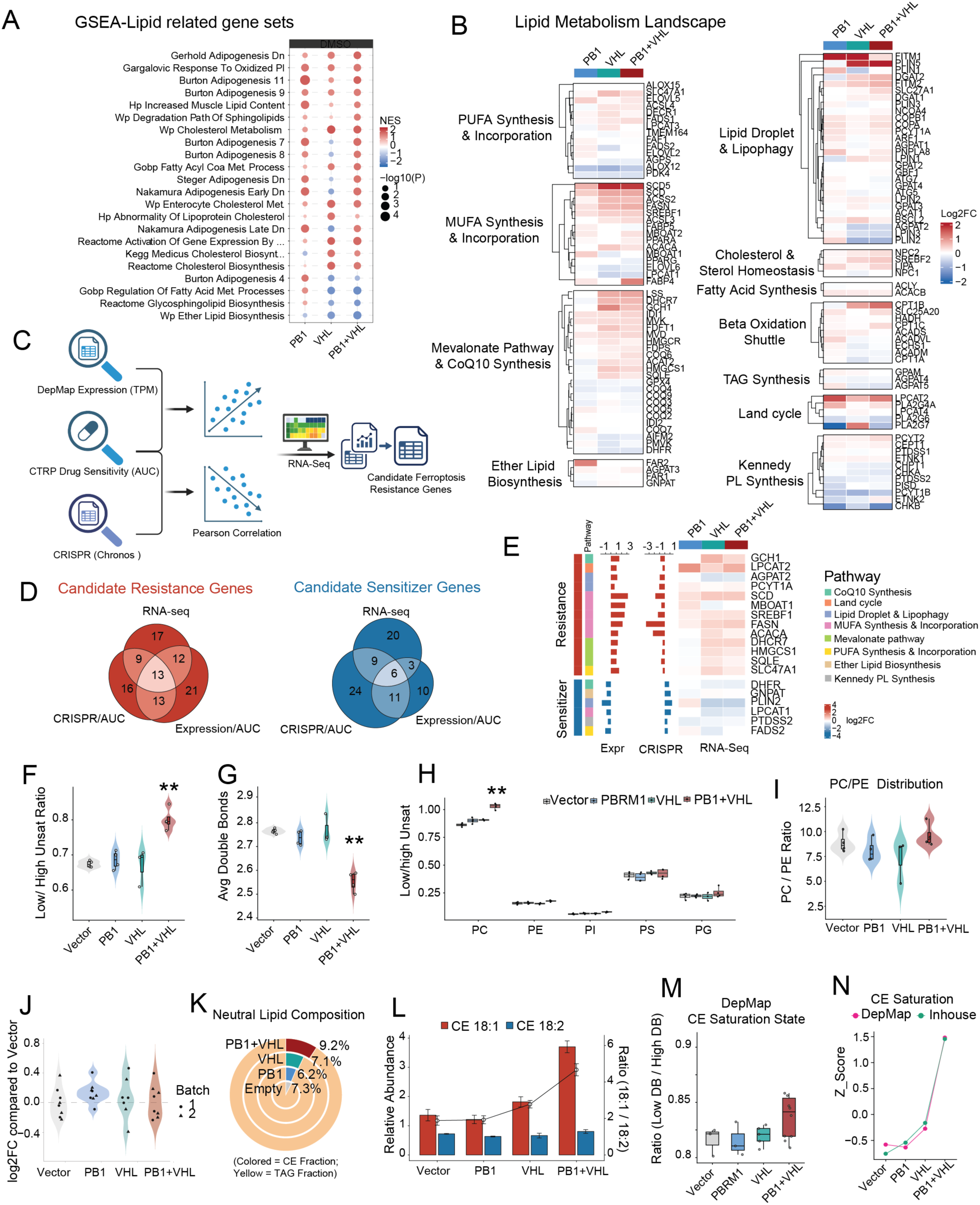
PBRM1 and VHL coordinately reprogram lipid metabolism to establish a ferroptosis-resistant transcriptional state. (A) GSEA bubble plot of lipid metabolism-related gene sets from the MSigDB C2 curated and C5 ontology collections in PBRM1, VHL, and PBRM1+VHL reconstituted Caki-2 cells relative to Empty vector under DMSO conditions at T2. Bubble size represents −log10(*P*adj) and fill color represents NES. Only gene sets with adjusted *P* < 0.05 are shown. (B) Heatmap of log2 fold-changes for a curated panel of 119 lipid metabolism genes organized by functional category in PBRM1, VHL, and PBRM1+VHL reconstituted cells relative to Empty vector under DMSO conditions at T2. Color scale represents log2 fold-change relative to Empty vector. (C) Schematic of the integrative DepMap pharmacogenomic analysis pipeline. Basal gene expression (TPM) and ferroptosis inducer drug sensitivity (CTRP AUC; RSL3, ML162, ML210, erastin) were correlated by Pearson correlation for lipid metabolism differentially expressed genes and Z-scored across the full compound library. CRISPR gene effect scores (Chronos) were separately correlated with GPX4 inhibitor AUC and *Z*-scored across all tested genes. Both scores were used to prioritize candidate ferroptosis resistance and sensitizer genes. (D) Venn diagrams showing the intersection of three evidence layers for candidate resistance genes (left, red) and candidate sensitizer genes (right, blue): RNA-seq differential expression in reconstituted cells, expression-AUC pharmacogenomic correlation, and CRISPR dependency-AUC correlation. Numbers indicate genes falling within each overlap region. (E) Integrated summary of validated dual candidates meeting both expression and CRISPR thresholds. Left bars show the expression–AUC correlation *Z*-score (averaged across RSL3, ML162, ML210, erastin); middle bars show the CRISPR dependency–AUC correlation *Z*-score (averaged across RSL3, ML162, ML210); right heatmap shows log2 fold-changes from RNA-seq at T2. Genes are grouped into resistance (upper) and sensitizer (lower) categories and annotated by metabolic pathway. (F) Violin plots of the global low-to-high unsaturation lipid species ratio (double bonds ≤ 1 versus ≥ 2) across Empty vector, PBRM1, VHL, and PBRM1+VHL Caki-2 cells measured by untargeted lipidomic. Higher ratios indicate a more MUFA-enriched membrane lipid composition. (G) Violin plots of the average number of double bonds per phospholipid acyl chain across genotypes, as a continuous membrane unsaturation index. Lower values reflect a less peroxidation-prone membrane composition. (H) Low-to-high unsaturation ratio as in (F), resolved by phospholipid class (PC, PE, PI, PS, PG) across all four genotypes. (I) Violin plots of the PC/PE molar ratio across genotypes. An elevated PC/PE ratio in reconstituted cells is consistent with a relative reduction of PE species, which constitute the proximal substrates of ferroptotic lipid peroxidation. (J) Log2 fold-changes of individual lipid species in PBRM1, VHL, and PBRM1+VHL cells relative to Empty vector, shown across two independent lipidomics batches. Lipid abundances were log2-transformed and normalized within each batch by subtracting the mean log2 value of Empty vector samples, centering Empty at zero while preserving biological variability. Individual biological replicates are shown with batch identity indicated by point shape; genotype means are shown as mean ± SEM. (K) Donut charts of neutral lipid class composition for each genotype. Darker/teal segments represent the cholesteryl ester (CE) fraction; yellow/pale segments represent the triacylglycerol (TAG) fraction. The percentage of CE within total neutral lipids is indicated for each genotype. (L) Relative abundance of CE 18:1 and CE 18:2 species (left y-axis, bars) and the CE 18:1/CE 18:2 ratio (right y-axis, line) across Empty vector, PBRM1, VHL, and PBRM1+VHL cells. Elevated CE 18:1 relative to CE 18:2 in reconstituted cells reflects preferential accumulation of oleate-esterified cholesteryl species and a reduction in oxidizable linoleate-containing neutral lipids. (M) CE saturation state across renal carcinoma cell lines in DepMap, stratified by genotype class as in Figure 3A. CE saturation was calculated as the ratio of low-unsaturation to high-unsaturation CE species. Box plots display the median with interquartile range. (N) Cross-dataset validation of CE saturation state. CE saturation Z-scores derived from DepMap kidney cancer cell line lipidomics and in-house Caki-2 lipidomics are plotted side by side across genotypes. Concordance between datasets supports the generalizability of the CE 18:1-enriched neutral lipid phenotype in PBRM1/VHL-reconstituted cells. (F, G, I) Lipidomics data are from batch 2 (*n* = 4 biological replicates per genotype, except VHL, *n* = 3; see Methods). Statistical significance was assessed by unpaired two-sided Welch’s *t*-test relative to Empty vector. (H) Lipidomics data from batch 2 as in (F, G, I); Welch’s *t*-test within each phospholipid class relative to Empty vector. (J) Data from both independent lipidomics batches (batch 1: *n* = 4 per genotype; batch 2: *n* = 4 per genotype except VHL, *n* = 3). Batch identity is shown by point shape; genotype means are mean ± SEM. Statistical comparisons were performed within each batch using Welch’s *t*-test on log₂-transformed values; batch-specific log₂ fold-change estimates were then combined using fixed-effect meta-analysis. (K, L) Lipidomics data from batch 2 as in (F, G, I). (L) Data are mean ± SEM; statistical significance was assessed by unpaired two-sided Welch’s *t*-test relative to Empty vector. (M, N) DepMap kidney cancer cell-line lipidomics data (publicly available); no new experimental replicates. \**P* ≤ 0.05; \*\**P* ≤ 0.01; \*\*\**P* ≤ 0.001; \*\*\*\**P* ≤ 0.0001.

### PBRM1 and VHL promote ferroptosis-resistant lipid remodeling of membranes

Lipidomic profiling confirmed global membrane remodeling. The dual PBRM1+VHL state showed enrichment for low-unsaturation lipid species and reduced average acyl-chain double bonds (Figs. 7F, G). This effect was most pronounced in phosphatidylcholine, the most abundant lipid class (Fig 7H), with species-level enrichment following these trends (S4I, J). The dual PBRM1+VHL re-expression was also accompanied by an increased PC/PE ratio (Figs. 7I), consistent with depletion of PUFA-PE ferroptotic substrates (23,24). Neutral lipid content was preserved overall (Fig 7J), but cholesterol esters were enriched (Fig. 7K), as a result of a selective increase in monounsaturated CE species (Figs. 7L, S4K). These lipidomic signatures were recapitulated in DepMap kidney cancer cell lines expressing PBRM1 and VHL (Figs. 7M, N). Together, these results demonstrate that PBRM1 and VHL cooperatively enforce a ferroptosis-resistant lipid state characterized by reduced membrane oxidizability and restricted availability of peroxidation substrates.

## Discussion

The data presented here identify PBRM1 as a modulator of ferroptotic cell death and demonstrate that VHL and PBRM1 cooperatively restrict GPX4 inhibitor sensitivity in ccRCC through largely independent transcriptional programs. Given the high frequency of PBRM1 loss in ccRCC, these findings link chromatin regulation directly to ferroptotic vulnerability in a clinically relevant tumor context.

The intrinsic GPX4 dependency of ccRCC reflects the metabolic consequences of VHL loss, which stabilizes HIF1A and HIF2α. Prior work has established HIF2α as a central driver of ferroptosis susceptibility in cancers by promoting PUFA enrichment in lipid droplets (8), and potentiating iron-dependent oxidative cell death (25). In parallel, HIF1A increases cellular iron availability through transcriptional upregulation of the transferrin receptor TFRC(26) and heme oxygenase-1 (HMOX1) (27). HIF1A depletion reduced RSL3 sensitivity in VHL-null Caki-2 cells. Consistent with this, VHL re-expression suppressed lipid peroxidation and GPX4 inhibitor sensitivity across multiple ccRCC models, in agreement with prior work showing that VHL represses HIF-driven ferroptotic vulnerability (7).

While loss of VHL is the primary determinant of ferroptosis sensitivity in ccRCC (7), we find that loss of PBRM1 confers additional sensitivity, similar to SETD2 (19), and BAP1 (20). Multiple SWI/SNF subunits have been implicated in ferroptosis through distinct mechanisms (21,28–32) , but this is the first study to specifically define a role for PBRM1. Although PBRM1’s function as a tumor suppressor in ccRCC cell lines is dependent on HIF1A (33), PBRM1-mediated ferroptosis resistance is not; genome-wide transcriptional programs regulated by PBRM1 and VHL were largely non-overlapping and additive. This distinguishes PBRM1 from both VHL-dependent HIF regulation and other ccRCC-mutated epigenetic regulators that sensitize cells to ferroptosis through iron accumulation (34) or cystine deprivation (20).

Our time-resolved transcriptomic profiling supports a ground-state resistance model. This is consistent with the established biochemistry of ferroptosis initiation: GPX4 inhibition triggers lipid peroxidation on a timescale too rapid for transcriptional induction (1,21,35). Therefore, any transcriptional changes observed following RSL3 treatment reflect a response to lipid peroxide accumulation rather than the cause. In contrast, a lack of transcriptional changes after RSL3 treatment in resistant cells reflects the pre-existing transcriptional programs at the ground state that govern the threshold for ferroptotic execution(23,24,36). Despite both conferring protection to ferroptosis, VHL and PBRM1 regulate distinct pathways. VHL re-expression uniquely decreases hypoxia, glycolysis and mTOR signaling, which are canonical HIF-driven processes linked to ferroptosis (8,26,37,38), while PBRM1 re-expression uniquely induced a partial EMT-like program enriched for extracellular matrix and collagen genes, that may confer resistance through altered membrane mechanics (39–42).

Focused analysis identified two major protective pathways shared by VHL and PBRM1. First, iron metabolism was remodeled toward sequestration and efflux with upregulation of *FTH1*, *FTL*, and the iron exporter *SLC40A1*, and repression of iron importers *TFRC* and *SLC11A2*, resulting in reduced intracellular Fe²⁺. Second, extensive lipid metabolic preprogramming reduced the oxidizability of membranes. Integration with DepMap data identified a core resistance signature spanning distinct metabolic pathways. The first is MUFA synthesis (*SREBF1*, *ACACA*, *FASN*, *SCD*), which competitively displaces oxidation-prone PUFAs from membranes. The second is phospholipid remodeling (*PCYT1A*, *LPCAT2*, *AGPAT2*, *MBOAT1*), which collectively routes MUFA-enriched acyl chains into membrane phospholipids. The third is sterol metabolism (*HMGCS1*, *SQLE*, *DHCR7*), which enriches cholesterol in membranes.

Among resistance-associated genes, SCD emerged as a key convergent node and pharmacological inhibition of SCD significantly attenuated the ferroptosis resistance conferred by VHL/PBRM1 re-expression, supporting MUFA biosynthesis as a critical part of VHL/PBRM1-mediated ferroptosis resistance. In parallel, lipid metabolism genes that sensitize cells to ferroptosis were downregulated across all reconstituted genotypes, including *PLIN2*, which coats PUFA-containing lipid droplets that can release oxidizable acyl chains upon lipophagy-driven turnover (43,44), *FADS2*, which is required for PUFA biosynthesis (45–47), and *GNPAT*, which is required for ether-PUFA-PE species (48). Together, these changes add a second dimension to the resistance phenotype: beyond upregulating MUFA biosynthesis, PBRM1 and VHL reduce the production of PUFA substrates for peroxidation.

Lipidomic analyses confirmed this shift toward MUFA-enriched, low-unsaturation membrane phospholipids, elevated PC/PE ratio and depletion of PUFA-PE substrates for peroxidation (23,24,49). The increase in MUFAs suppresses ferroptosis by competing with PUFAs for membrane phospholipid incorporation (50), while an increase in low-unsaturation cholesterol esters limits the availability of oxidizable PUFA acyl chains (51–53). ER-mitochondria contact sites are intracellular hotspots of ferroptotic phospholipid peroxidation and are enriched in di-PUFA-PE and di-PUFA-PC species (35). This suggests that the depletion of these doubly unsaturated species may reduce the peroxidation-prone substrate pool at the precise subcellular compartment where ferroptotic lipid oxidation is initiated.

Several limitations warrant consideration. The functional relevance of the PBRM1-associated partial EMT program has not been tested and whether these mechanisms operate *in vivo* remains to be determined. Nonetheless, co-loss of VHL and PBRM1 occurs in approximately 40% of ccRCC tumors, a cancer lineage that shows pronounced GPX4 dependency. These findings nominate PBRM1 status as a potential modifier of ferroptosis sensitivity and suggest that chromatin-regulated iron and lipid programs may represent intrinsic resistance mechanisms with relevance to ferroptosis-based therapeutic strategies.

## Acknowledgements

ECD is supported by NIH U01CA207532, V Foundation for Cancer Research (V2014-004 and D2016-030), American Cancer Society (RSG-21-012-01-DMC) and Indiana CTSI (TR001108). GJ is supported by a SIRG graduate fellowship from the Purdue Institute for Cancer Research. We acknowledge support from the IU Simon Cancer Center (Grant P30CA082709), Purdue University Center for Cancer Research (Grant P30CA023168) and Walther Cancer Foundation. We acknowledge the Purdue Genomics Core and the Purdue Cytometry and Cell Separation core for technical support and the Purdue University’s Metabolite Profiling Facility in the acquisition and analysis of mass spectrometry data. We thank Qing Zhang (UTSW) for the pBABE-VHL plasmid and RCC4 cell line, and Walter Storkus (U. Pitt) for the SLR25 cell line.

## Conflict of Interest

The authors have no conflicts of interest to declare.

## Availability of Data and Materials

All plasmids have been deposited at Addgene. RNA-seq data generated in this study have been deposited in the NCBI Gene Expression Omnibus (GEO) under accession number GSEXXXXXX. CUT&RUN sequencing data have been deposited under accession number GSEXXXXXX. Lipidomics data are available as Supplementary Data and the raw data is deposited in Purdue University Research Repository.

All other data supporting the conclusions of this study are included in the supplementary files or available from the corresponding author upon reasonable request.

## Materials and Methods

### Cell lines and culture conditions

Caki-2, HEK293T, MDA-MB-231, BT-549, A172, LNCaP, and 22Rv1 cells, were obtained from the American Type Culture Collection (ATCC, Manassas, VA, USA). RCC4 was a gift from Qing Zhang (UTSW) and SLR25 was a gift from Walter Storkus (U. Pitt. SoM). Caki-2 cells were cultured in McCoy’s 5A medium (Corning Mediatech, Corning, NY, USA) supplemented with 10% fetal bovine serum (FBS) (Corning Mediatech, Corning, NY, USA ), 100 U/mL penicillin, 100 µg/mL streptomycin (Corning Mediatech, Corning, NY, USA ), and 2 mM L-alanyl-L-glutamine (GlutaGro™, Corning Mediatech, Corning, NY, USA ). RCC4, SLR25, A172, MDA-MB-231, and HEK293T cells were maintained in high-glucose DMEM (Corning Mediatech, Corning, NY, USA) with 10% FBS and 1% penicillin-streptomycin. BT-549 cells were cultured in RPMI-1640 (Corning Mediatech, Corning, NY, USA) with 10% FBS, 1% penicillin-streptomycin, and 0.023 U/mL insulin. LNCaP and 22Rv1 cells were maintained in phenol-red-free RPMI-1640 with 10% FBS, 100 U/mL penicillin, 100 µg/mL streptomycin, and 2 mM L-alanyl-L-glutamine.

All cell lines were cultured at 37 °C in a humidified 5% CO₂ incubator. Plasmocin™ (InvivoGen, San Diego, CA, USA; 1:10 000 dilution) was added routinely to prevent mycoplasma contamination, and all lines were periodically confirmed mycoplasma-free. Cells were passaged at approximately 80% confluence using 0.05% trypsin-EDTA, and media were replaced every 2-3 day.

### Reagents

RSL3 ((1S,3R)-RSL3, Selleckchem, Houston, TX, USA; S8155), ML162 (Selleckchem, Houston, TX, USA, S4452), imidazole ketone erastin (IKE; Selleckchem, Houston, TX, USA, S8877), ferrostatin-1 (Fer-1; Selleckchem, Houston, TX, USA, S7243), and CAY10566 (Cayman Chemical, Ann Arbor, MI, USA, 10012562) were dissolved in DMSO and stored at −20 °C. Ammonium iron(II) sulfate hexahydrate (Sigma-Aldrich, St. Louis, MO, USA; 09719) and deferoxamine mesylate (DFO; Cayman Chemical, Ann Arbor, MI, USA, 14595) were dissolved in sterile water or PBS and prepared freshly before each use. DMSO (vehicle control) was used at a final concentration not exceeding 0.1% (v/v) in all assays.

### Antibodies

The following primary antibodies were used for immunoblotting at 1:1000 dilution unless otherwise indicated: anti-PBRM1/BAF180 (Abcam, Cambridge, UK; ab243876); anti-PHF10/BAF45A (Invitrogen, Waltham, MA, USA, PA5-30678); anti-VHL (Santa Cruz Biotechnology, Dallas, TX, USA; sc-135657); anti-GPX4 (Santa Cruz Biotechnology, sc-166570); anti-FTH1 (Santa Cruz Biotechnology, sc-376594); anti-FTL (Cell Signaling Technology, Danvers, MA, USA; 68106T); anti-TFRC (Santa Cruz Biotechnology, sc-65882); anti-HIF1A (Cell Signaling Technology, 36169); anti-β-actin (Santa Cruz Biotechnology, sc-47778); and anti-PCNA (Santa Cruz Biotechnology, sc-56). For CUT&RUN, the following antibodies were used: anti-PBRM1/BAF180 (Bethyl Laboratories, Montgomery, TX, USA; A301-591A); anti-PHF10/BAF45A (Invitrogen, Waltham, MA, USA, PA5-30678); anti-H3K4me3 (EpiCypher, Durham, NC, USA; 13-0060); and rabbit IgG (EpiCypher, 13-0042) as a negative control. IRDye 800CW- and IRDye 680RD-conjugated secondary antibodies (LI-COR Biotechnology, Lincoln, NE, USA) were used at 1:10000 for all immunoblotting.

### Stable cell line generation and lentiviral transduction

Lentivirus was produced in HEK293T cells by co-transfection of lentiviral expression constructs with the packaging plasmids pMD2.G and psPAX2. Viral supernatants were collected 48 h post-transfection and concentrated by ultracentrifugation at 17,500 rpm (∼52,000 × g, Beckman Coulter SW 32 Ti rotor, Brea, CA, USA) for 2 h. The viral pellet was resuspended in 200 µL PBS. Caki-2, SLR25, and HEK293T cells were transduced with concentrated lentivirus by spinfection (1,500 rpm, ∼250 × g, 1 h, room temperature, swinging-bucket rotor). Cells were incubated for 48 h post-infection, then selected with puromycin (2 µg/mL; Sigma-Aldrich, St. Louis, MO, USA) and hygromycin (200 µg/mL; Corning Mediatech, Corning, NY, USA) where applicable. The efficiency of all constructs was confirmed by immunoblotting.

Stable re-expression of PBRM1 and VHL was achieved using a doxycycline-inducible lentiviral system. Full-length PBRM1 was subcloned from pBabe-puro-BAF180 (gift from Ramon Parsons; Addgene, Watertown, MA, USA; #41078) into the tetracycline-inducible vector TetO-FUW (gift from Rudolf Jaenisch; Addgene, #20323) carrying a puromycin resistance cassette. Full-length VHL was cloned from pBABEhygro VHL-FLAG (gift from Qing Zhang, UTSW) into a modified TetO-FUW carrying a blasticidin resistance cassette. Inducible transgene expression in all lines was driven by the reverse tetracycline transactivator encoded in pLenti-CMV-rtTA3-Hygro (w785-1; gift from Eric Campeau; Addgene, #26730).

To generate isogenic lines for direct comparison, all cells were co-transduced with both a puromycin-resistant and a blasticidin-resistant TetO-FUW construct as follows: PBRM1-reconstituted cells received TetO-FUW-PBRM1-puro and empty TetO-FUW-blast; VHL-reconstituted cells received TetO-FUW-VHL-blast and empty TetO-FUW-puro; PBRM1+VHL double-reconstituted cells received TetO-FUW-PBRM1-puro and TetO-FUW-VHL-blast; Empty vector controls received both empty constructs. This design ensures that all four genotype groups carry equivalent numbers of integrated constructs and are maintained under identical dual selection pressure. Stable integrants were selected with hygromycin (200 µg/mL; Corning Mediatech, Corning, NY, USA), puromycin (2 µg/mL; Sigma-Aldrich, St. Louis, MO, USA), and blasticidin (10 µg/mL; Thermo Fisher Scientific, Waltham, MA, USA; A1113903) for 5 days.

Transgene expression was induced by addition of doxycycline (1 µg/mL) for three days prior to all downstream experiments. Re-expression was confirmed by immunoblotting before use.

### CRISPR-Cas9-mediated gene disruption

CRISPR-Cas9-mediated gene disruption was performed in HEK293T cells using sgRNA sequences cloned into lentiCRISPRv2 (Addgene, #52961; gift from Dr. Zhang Feng), which encodes Cas9 alongside either a puromycin or blasticidin resistance cassette (gift from Dr. Nathaniel Mabe). To achieve simultaneous disruption of PBRM1 and VHL, sgPBRM1 was cloned into the puromycin-resistant vector and sgVHL into the blasticidin-resistant vector. Guide sequences used were: sgPBRM1, 5’-TTCATCCTTATAGTCTCGGA-3’; sgVHL, 5’-CGCGCGTCGTGCTGCCCGTA-3’. A non-targeting control used the sgChr2-2 sequence in both vectors, and control cells were co-transduced and selected in parallel (sgChr2-2: 5’-GGTGTGCGTATGAAGCAGTG-3’). Stably transduced cells were selected with puromycin (2 µg/mL) and blasticidin (10 µg/mL) for 4 days. Gene disruption was confirmed by immunoblotting.

### shRNA-mediated gene knockdown

Inducible shRNA knockdown was performed using constructs cloned into pLKO-Tet-On-blasticidin, a modified version of pLKO-Tet-On in which the puromycin resistance cassette was replaced with blasticidin to permit knockdown in Caki-2 cells already under puromycin selection. The following sequences were used: shHIF1A-3809 (TRCN0000003809), 5’-CCAGTTATGATTGTGAAGTTA-3’; shHIF1A-3810 (TRCN0000003810), 5’-GTGATGAAAGAATTACCGAAT-3’; shHIF1A-0819 (TRCN0000010819), 5’-TGCTCTTTGTGGTTGGATCTA-3’. shPBRM1 (TRCN0000015994), 5’-TTTGTAGATCAAAGACTCCGG-3’ was used for BT549 with pLKO-Tet-On-puro. A non-targeting scramble shRNA (5’-GCTACACTATCGAGCAATT-3’) served as control in all experiments. Cells were selected with blasticidin (10 µg/mL) for two weeks. shRNA expression was induced with doxycycline (1 µg/mL) for 3 days before downstream experiments. Knockdown efficiency was confirmed by immunoblotting.

For HIF1A knockdown in Caki-2, cells were first transduced with TetO-FUW-PBRM1-puro (or empty TetO-FUW-puro and selected with puromycin (2 µg/mL), then transduced with the shHIF1A or shScramble constructs and selected with blasticidin (10 µg/mL) for one week. For BT549, cells were transduced directly with shPBRM1 or shScramble and selected with puromycin (2 µg/mL).

### Immunoblotting

Cells were harvested by trypsinization and lysed in RIPA buffer (50 mM Tris-HCl pH 8.0, 150 mM NaCl, 0.1% SDS, 0.5% sodium deoxycholate, 1% NP-40) with freshly added protease inhibitors (PMSF, aprotinin, leupeptin, and pepstatin) for 30 min on ice, then clarified by centrifugation at 21,000 × g for 15 min at 4 °C. For nuclear extracts, cells were resuspended at approximately 2 × 10⁷ cells/mL in Buffer A (20 mM HEPES pH 7.9, 25 mM KCl, 10% glycerol, 0.1% NP-40) with protease inhibitors and incubated on ice for 5 min. Nuclei were pelleted at 600 × g for 10 min, resuspended in chromatin IP buffer (20 mM HEPES pH 7.9, 150 mM NaCl, 1% Triton X-100, 7.5 mM MgCl₂, 0.1 mM CaCl₂) with protease inhibitors, and centrifuged at 21,000 × g for 30 min at 4 °C.

Protein concentrations were determined by BCA assay (Pierce Biotechnology, Rockford, IL, USA) with BSA as standard. Equal amounts of protein were combined with 4× LDS sample buffer containing 10% β-mercaptoethanol, denatured at 95 °C for 5 min, and resolved on 4-12% Bis-Tris gradient gels (Invitrogen, Waltham, MA, USA, NW04120BOX). Proteins were transferred to PVDF membranes, blocked in 5% BSA/TBS-T (0.1% Tween-20) for 30 min, and incubated with primary antibodies overnight at 4 °C. After washing, membranes were incubated with IRDye-conjugated secondary antibodies (LI-COR Biotechnology, Lincoln, NE, USA) for 1 h at room temperature. Signals were detected on an Odyssey CLx system (LI-COR Biotechnology, Lincoln, NE, USA) and quantified using Image Studio software. Full-length uncropped blots are provided as Supplementary material.

### Cell viability assays

Cell viability was assessed using the CCK-8 assay (MedChemExpress, Monmouth Junction, NJ, USA; HY-K0301) or CellTiter-Glo® Luminescent Cell Viability Assay (Promega, Madison, WI, USA, G7572) according to the manufacturers’ instructions. Cells were seeded at 5 000 per well in 96-well plates 24 h before treatment. For doxycycline-inducible lines, cells were pre-treated with doxycycline (1 µg/mL) for 72 h before compound addition to ensure full transgene induction. This step was omitted for constitutive HEK293T CRISPR-edited lines. Cells were then exposed to the indicated compounds for approximately 16-18 h. CCK-8 reagent was added to the culture medium and plates were incubated at 37 °C for 3 h before absorbance was read at 450 nm on a GloMax^®^ Discover Microplate Reader (Promega, Madison, WI, USA).

For SCD inhibitor experiments, cells were treated with CAY10566 (0.5 µM) for 24 h after seeding, followed by RSL3 (0.33 µM) overnight before viability assessment by CellTiter-Glo^®^. Viability was normalized to DMSO-treated controls within each experiment and genotype. IC₅₀ values were calculated by nonlinear regression using a four-parameter log-logistic model in GraphPad Prism 10 (GraphPad Software, San Diego, CA, USA).

### Flow cytometric assessment of lipid peroxidation

Lipid peroxidation was assessed using BODIPY™ 581/591 C11 (Invitrogen, Waltham, MA, USA, D3861) and Liperfluo (Dojindo, Kumamoto, Japan; L248). For BODIPY™ 581/591 C11, cells seeded at 70-80% confluence in 12- or 24-well plates were treated with ferroptosis inducers for 8-24 h, washed with serum-free medium, incubated with 5 µM probe for 30 min at 37 °C protected from light, washed with PBS, and harvested by trypsinization for immediate analysis. Data were acquired on an Attune NxT flow cytometer (Thermo Fisher Scientific, Waltham, MA, USA) using the BL1 (oxidized, green) and YL1 (reduced, red) channels; the BL1/YL1 fluorescence ratio served as a quantitative index of lipid peroxidation.

For Liperfluo staining, a 1 mM DMSO stock was prepared within 24 h of use. Cells were seeded and treated as above, then incubated with 1 µM Liperfluo in serum-free medium for 30 min at 37 °C protected from light, washed twice with PBS, and analyzed immediately on the Attune NxT using the BL1 channel. A minimum of 10 000 single-cell events were collected per sample for both probes. Mean fluorescence intensity (MFI) values were normalized to DMSO-treated Empty vector controls.

### Measurement of labile ferrous iron by FerroOrange

Intracellular labile Fe²⁺ was measured using FerroOrange (Cayman Chemical, Ann Arbor, MI, USA, 41725). A 1 mM stock in DMSO was stored at −20 °C protected from light, and a 1 µM working solution was prepared in PBS immediately before use. Cells grown to 70-80% confluence were washed three times with serum-free medium after the indicated treatment, incubated with 1 µM FerroOrange for 30 min at 37 °C protected from light, and harvested by trypsinization without further washing to prevent probe leakage. Cells were resuspended in PBS and analyzed immediately on the Attune NxT using the YL1 channel. A minimum of 10 000 events per sample were collected and MFI values normalized to the DMSO-treated Empty vector control.

### Colorimetric quantification of total cellular iron

Total cellular iron (Fe²⁺ + Fe³⁺) was quantified using the Iron Assay Kit (Abcam, ab83366). Cells were washed twice with cold PBS and lysed in iron assay buffer on ice. Samples were subjected to three freeze-thaw cycles (liquid nitrogen/37 °C) to ensure complete iron release, then centrifuged at 13,000 × g for 10 min at 4 °C. Iron probe reagent was added to clarified supernatants and incubated for 60 min at room temperature protected from light. Absorbance was measured at 593 nm on a GloMax® Discover Microplate Reader (Promega, Madison, WI, USA) and normalized to cell number determined by CellTiter-Glo® (Promega, Madison, WI, USA). Values were expressed relative to the Empty vector control.

### GSH/GSSG ratio measurement

The GSH/GSSG ratio was determined using the GSH/GSSG-Glo™ Assay (Promega, Madison, WI, USA, V6611) following the manufacturer’s instructions. Cells were seeded in white 96-well plates and treated with DMSO or RSL3 (0.33 µM) for 6 h. Luminescence was measured on a GloMax® Discover Microplate Reader (Promega, Madison, WI, USA). The GSH/GSSG ratio was calculated from individual luminescence values and expressed relative to the DMSO-treated Empty vector control.

### NAD⁺/NADH pool quantification

Total NAD⁺ + NADH was quantified using the NAD/NADH-Glo™ Assay (Promega, Madison, WI, USA, G9071). Cells were treated with DMSO or RSL3 (0.33 µM) for 6 h and processed per the kit protocol. Luminescence was measured on a GloMax® Discover Microplate Reader (Promega, Madison, WI, USA). Values were normalized to cell number and expressed relative to the Empty vector control.

### DepMap and CTRP pharmacogenomic analyses

CRISPR dependency scores (Chronos), baseline transcriptomic profiles (TPM), and lipidomics data were retrieved from the Broad Institute Cancer Dependency Map (DepMap, release 25Q3+; https://depmap.org/portal/) (54). Cell-line lineage and subtype annotations were obtained from DepMap metadata and the Cancer Cell Line Encyclopedia (CCLE) (55). Drug sensitivity profiles, quantified as area under the dose–response curve (AUC), were obtained from the Cancer Therapeutics Response Portal v2.0 (CTRPv2; https://portals.broadinstitute.org/ctrp.v2/) (56) and annotated using CCLE identifiers. All downstream analyses were performed in R (v4.4.1) using data.table (v1.15.4), tidyverse (v2.0.0), ggplot2 (v3.5.1), broom (v1.0.6), and boot (v1.3-28).

For GPX4 dependency and lineage analyses, GPX4 Chronos scores were extracted across all human cancer cell lines. Kidney-derived lines were classified as renal cell carcinoma (RCC) or non-RCC kidney, and subtypes (clear cell, papillary, unclassified) were distinguished within the RCC group. For lineage-level comparisons of GPX4 dependency and GPX4-inhibitor AUC (RSL3, ML162, ML210), median values were calculated per lineage and differences between kidney and non-kidney lineages assessed using two-sided Wilcoxon rank-sum tests.

For Genotype stratification, PBRM1/VHL loss status was defined by the presence of copy number loss, a damaging mutation, or a hotspot mutation in DepMap; wildtype status required all three criteria to be absent. Each renal carcinoma line was assigned to one of four groups: both wildtypes, VHL loss only, PBRM1 loss only, or dual loss. Effect sizes for the association between genotype and GPX4 dependency were estimated as point-biserial Pearson correlations with 95% confidence intervals from 1,000-iteration bootstrap resampling (percentile method).

For Dual-omics prioritization of candidate lipid genes, to prioritize lipid metabolism DEGs coupled to ferroptosis outcomes, a dual-omics correlational framework was applied. In the expression arm, Pearson correlation coefficients between basal gene expression and drug AUC were computed across all overlapping cell lines, then Z-scored across the full CTRP drug library (∼481 compounds) for each gene; this normalization captures ferroptosis-specific expression–sensitivity associations above and beyond general drug response. A positive expression Z-score (*Z*_expr_ > 0.5) indicates that elevated expression associates with ferroptosis resistance; a negative score (*Z*_expr_ < −0.5) indicates association with sensitivity. In the CRISPR arm, gene effect scores were correlated with mean AUC of three GPX4 inhibitors (RSL3, ML162, ML210) and Z-scored across all tested genes. A negative CRISPR Z-score (*Z*_CRISPR_ < −0.3) identifies genes disproportionately essential in resistant lines, reflecting functional dependence of the resistant state on those genes; a positive score (*Z*_CRISPR_> 0.3) identifies genes more essential in sensitive lines. Resistance candidates were defined by the intersection of *Z*_expr_> 0.5 and *Z*_CRISPR_ < −0.3; sensitizer candidates by *Z*_expr_< −0.5 and *Z*_CRISPR_ > 0.3.

For DepMap lipidomics validation, cell-line lipidomics data were obtained from DepMap (57).Kidney cancer cell lines were stratified by PBRM1/VHL loss status using the criteria described above. For each cell line, the cholesteryl ester (CE) saturation state was calculated as the ratio of low-unsaturation (≤1 double bond) to high-unsaturation (≥2 double bonds) CE species. Z-scores were computed across genotype groups to enable cross-dataset comparison with in-house Caki-2 lipidomic. Box plots display the median with interquartile range.

### RNA isolation and sequencing

Caki-2 cells expressing Empty vector, PBRM1, VHL, or PBRM1+VHL were seeded in 6-well plates in three biological replicates and treated with DMSO or RSL3 (0.33 µM) for 4 h (T1) or 10 h (T2). Cells were harvested in TRIzol™ (Thermo Fisher Scientific, Waltham, MA, USA) and RNA was isolated using the PureLink™ RNA Isolation Kit (Invitrogen, Waltham, MA, USA, 12183018A). RNA quality was assessed by Agilent Bioanalyzer (Agilent Technologies, Santa Clara, CA, USA); samples with RNA Integrity Number (RIN) ≥ 9 were used for library preparation. Poly(A)-selected libraries were prepared with the Illumina Stranded mRNA Prep Ligation Kit (Illumina, San Diego, CA, USA; 20040534) and quality-checked by Qubit (Thermo Fisher Scientific, Waltham, MA, USA ) and Agilent Bioanalyzer. Libraries were sequenced with 150 bp paired-end reads on an Illumina NovaSeq X Plus (T1) at Indiana University Genomics Core Facility or on an Element AVITI (Element Biosciences, San Diego, CA, USA; T2) at Purdue Genomics Core Facility.

### RNA-seq data processing and differential expression analysis

Raw reads were trimmed using fastp (v0.24.0) (58) and aligned to the human reference genome (GRCh38) using STAR (v2.7.11a) (59)with default parameters. Gene-level counts were quantified with featureCounts from the Subread package (v2.0.6) (60) against Ensembl annotation release 110. Differential expression analysis was performed using DESeq2 (v1.42.0) in R (v4.4.1) (61). Each reconstituted genotype was compared against the time-matched Empty vector control within each treatment condition. Adjusted *P*-values were calculated using the Benjamini-Hochberg method; genes with adjusted *P* < 0.05 were considered differentially expressed. Volcano plots, heatmaps, and summary tables were generated in R.

### Combinatorial epistasis modeling

To evaluate the transcriptional interaction between PBRM1 and VHL, an additive model was applied to log₂ fold-change values. The predicted PBRM1+VHL profile was calculated as the mean of individual PBRM1 and VHL log₂ fold-changes relative to the time-matched Empty vector: Predicted log₂FC(PBRM1+VHL) = Mean(log₂FC(PBRM1), log₂FC(VHL)). Predicted values were correlated with observed values by Pearson correlation. Deviation of mean residuals (observed minus predicted) from zero was assessed using an unpaired two-sided Student’s t-test and a Wilcoxon signed-rank test. Orthogonality of single-genotype transcriptional programs was assessed by direct Pearson correlation of PBRM1 and VHL log₂ fold-change values. Points were classified as "additive" (|observed − predicted| < 1), "synergy" (observed − predicted > 1), or "buffering" (observed − predicted < −1).

### Gene set enrichment and over-representation analyses

GSEA was performed using fgsea (v1.28.0) (62) in R. Genes were pre-ranked by the product of the sign of log₂FC and −log₁₀(adjusted P-value). Gene sets from MSigDB v2023.2 (https://www.gsea-msigdb.org/gsea/msigdb) including Hallmark (H), C2 curated, and C5 ontology collections were used (63); sets with fewer than 15 or more than 500 members were excluded. Significance was defined as adjusted *P* < 0.05 (Benjamini-Hochberg). Over-representation analysis was performed using Enrichr (https://maayanlab.cloud/Enrichr/) (64)against KEGG 2021 Human and WikiPathways 2019 Human databases. The combined enrichment score (c = ln(p) × z) was used to scale bubble size in dot plots. Leading-edge heatmaps were generated using ComplexHeatmap (v2.18.0) in R (65).

### CUT&RUN chromatin profiling

CUT&RUN was performed in Caki-2 PBRM1-reconstituted cells following the EpiCypher protocol (v2.2) with minor modifications. In-house purified pAG-MNase was used; the construct was obtained from Addgene (Watertown, MA, USA; #123461; gift from Steven Henikoff) and purified as previously described. The amount used was 37 ng per reaction, and incubation with bead-bound cells was performed for 1 h at 4 °C. Antibodies were: anti-PBRM1 (Bethyl Laboratories, Montgomery, TX, USA; A301-591A), anti-PHF10 (Invitrogen, Waltham, MA, USA, PA5-30678), anti-H3K4me3 (EpiCypher, Durham, NC, USA; 13-0060), and rabbit IgG (EpiCypher, Durham, NC, USA; 13-0042). Libraries were prepared using the CUTANA™ CUT&RUN Library Prep Kit (EpiCypher, Durham, NC, USA; 14-1001 and 14-1002) and quality-checked by Qubit and Agilent TapeStation at Purdue Genomics Core Facility (Purdue University, West Lafayette, IN, USA). Libraries were sequenced with 150 bp paired-end reads on an Element AVITI at Purdue Genomics Core Facility (Purdue University, West Lafayette, IN, USA). All experiments were performed with two biological replicates.

### CUT&RUN data analysis

Raw paired-end reads were aligned using a dual-mapping strategy. Alignment to GRCh37/hg19 was performed with Bowtie2 (v2.5.4) (66) using local alignment parameters suited to size-selected CUT&RUN fragments (--local --very-sensitive-local --no-unal --no-mixed --no-discordant --phred33 -I 10 -X 700). Alignment to the E. coli K12 MG1655 spike-in genome was performed in a separate run using end-to-end parameters (--end-to-end --very-sensitive --no-overlap --no-dovetail --no-unal --no-mixed --no-discordant --phred33 -I 10 -X 700). BAM files were filtered with SAMtools (v1.21) (67)to retain properly paired, uniquely mapped reads (-f 2 -q 10). PCR duplicates were flagged using SAMtools markdup to assess library complexity but were not removed. Reads overlapping ENCODE GRCh37/hg19 blacklisted regions (v2) were excluded using BEDTools (v2.31.1) (68).

A spike-in scaling factor was calculated as 100 000 divided by the number of high-quality E. coli read pairs per sample, providing sequencing depth-independent normalization. Spike-in normalized bigWig tracks were generated using bamCoverage from deepTools (v3.5.4; --normalizeUsing None --scaleFactor [sample-specific value] --extendReads --exactScaling --binSize 10) (69). Genome browser tracks were visualized using IGV, displaying spike-in normalized bigWig files at the indicated loci with y-axes set to equivalent scales across antibodies All genomic coordinates are reported relative to GRCh37/hg19 .

### Lipidomic profiling and data analysis

Lipids were extracted following the method of Bligh and Dyer (70). Cell pellets were resuspended in 200 µL ultrapure water by vortexing. Methanol (550 µL) and HPLC-grade chloroform (250 µL) were added and the mixture vortexed for 10 s to produce a monophasic solution, then incubated at 4 °C for 15 min. Phase separation was induced by addition of ultrapure water (250 µL) and chloroform (250 µL). Samples were centrifuged at 16,000 × g for 10 min. The lower organic phase was carefully transferred to a clean microtube, avoiding the protein interphase. Solvent was evaporated under a nitrogen stream or in a SpeedVac. Dried extracts were stored at −80 °C until analysis; all samples were processed within two weeks of extraction to minimize lipid degradation.

Lipidomic profiling was performed by multiple reaction monitoring (MRM) profiling method, which includes flow-injection electrospray mass spectrometry without chromatographic separation(71). Dried extracts were reconstituted in injection solvent (acetonitrile/methanol/300 mM ammonium acetate, 3:6.65:0.35 v/v/v) and diluted into injection solvent spiked with 0.1 ng/µL Equisplash Lipidomics internal standard mix (Avanti Polar Lipids, Birmingham, AL, USA; 330731). An 8 µL volume was injected via micro-autosampler (Agilent G1377A) at 10 µL/min into the ESI source of an Agilent 6410 triple quadrupole mass spectrometer (Agilent Technologies). The capillary pump was operated at 150 bar, capillary voltage was 5 kV, and drying gas was maintained at 5.1 L/min at 300 °C. The MRM profiling method profiled multiple lipid classes including phospholipids, diacylglycerols, triacylglycerols, and cholesteryl esters, resolving differences in acyl chain length and unsaturation. For triacylglycerols, ammonium adducts were used as precursor ions and product ions represented neutral losses of selected fatty acyl residues (72). Raw MRM profiling data were processed using an in-house script to generate summed absolute ion intensities per MRM transition. All lipid abundances were normalized to internal standards.

Lipidomic profiling was performed in two independent experimental batches. Batch 1 included four biological replicates per genotype (Empty vector, PBRM1, VHL, PBRM1+VHL). Batch 2 included four biological replicates per genotype, except VHL, for which one replicate was excluded from downstream analysis due to low total lipid signal, yielding final sample sizes of *n* = 4 for Empty vector, PBRM1, and PBRM1+VHL, and *n* = 3 for VHL in batch 2. Exclusion was applied before statistical analysis based on pre-specified QC criteria. Because batch 2 provided more stable and broader lipid species coverage, all analyses shown in Figure 7 and Figure S4 use batch 2 data unless otherwise noted. For Figure 7J specifically, data from both batches were integrated to increase statistical power; batch-specific normalization was applied (lipid abundances log₂-transformed and centered by subtracting the mean of Empty vector samples within each batch), and batch-specific log₂ fold-change estimates were combined using fixed-effect meta-analysis. Lipid species were classified as low-unsaturation (≤1 double bond) or high-unsaturation (≥2 double bonds) based on sum-composition annotations. Statistical comparisons used Welch’s *t*-test on log₂-transformed values. Analyses were performed in R (v4.4.1).

### Statistical analysis

Statistical analyses were performed in GraphPad Prism 10 and R (v4.4.1). Data are presented as mean ± SD from n = 3 biologically independent replicates with individual data points shown, unless otherwise stated. Comparisons between two groups used an unpaired two-sided Student’s t-test. Comparisons among three or more groups used one-way ANOVA with Dunnett’s multiple-comparisons test with Empty vector as the reference. Lineage-level comparisons used two-sided Wilcoxon rank-sum tests. Exact *P*-values are reported where differences did not reach the threshold for asterisk annotation. For lipidomic comparisons, Welch’s *t*-test (unequal variance) was applied to log₂-transformed values to accommodate unequal sample sizes across genotypes. Significance thresholds: \**P* ≤ 0.05; **P ≤ 0.01; ***P ≤ 0.001; ****P ≤ 0.0001.

### Use of AI-assisted writing tools

Portions of this manuscript were edited with assistance from an artificial intelligence language model (Claude, Anthropic) and subsequently reviewed, revised, and verified for accuracy by the authors. All scientific content, analysis, data interpretation, and conclusions were performed by the authors.

---

**Figure S1.**
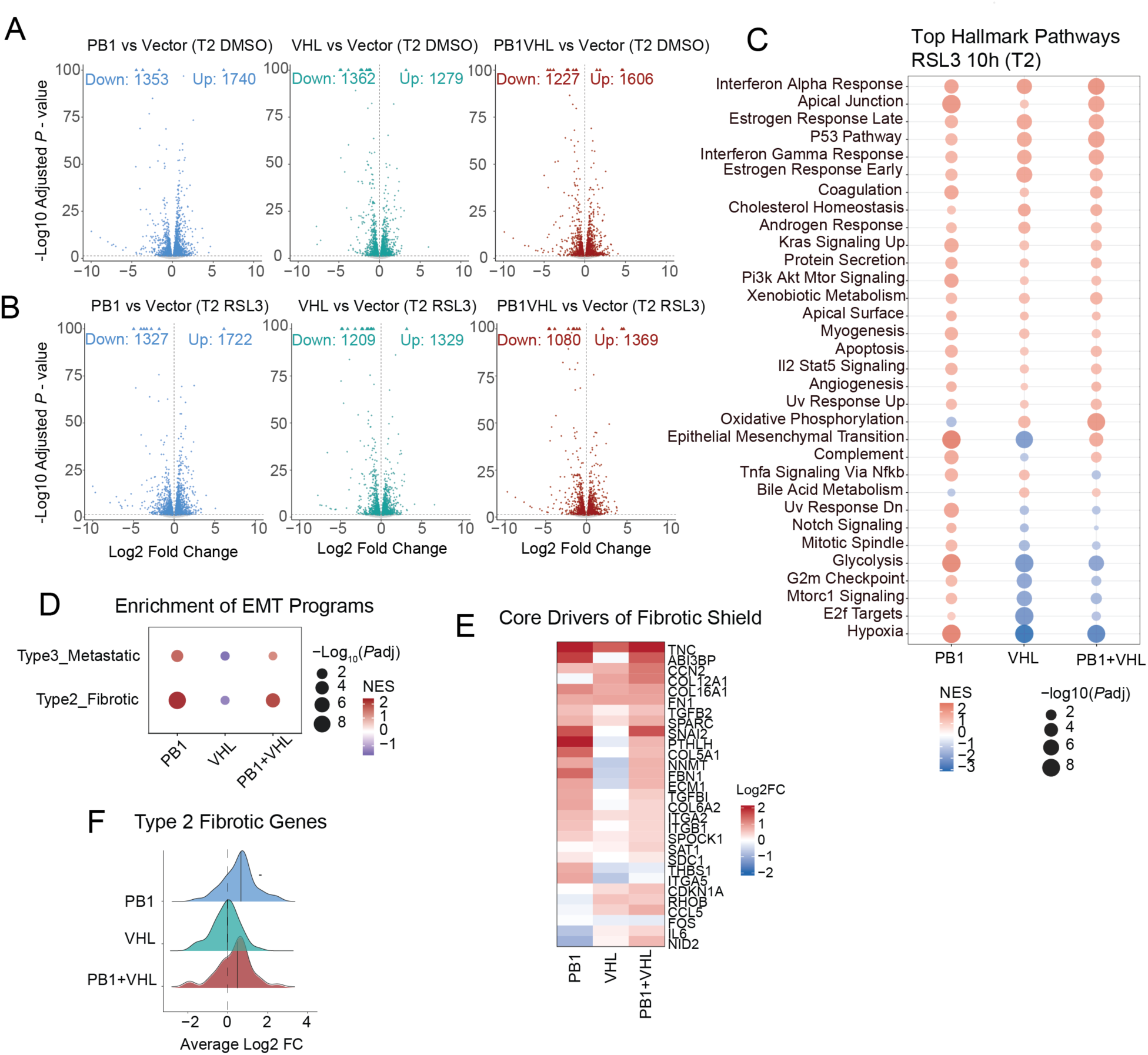
Validation of PBRM1 and VHL re-expression in additional ccRCC models, and PBRM1 depletion sensitizes BT549 cells to ferroptosis. (A) Immunoblot validation of PBRM1 and/or VHL re-expression in RCC4 cells. β-actin is shown as a loading control. (B) Immunoblot validation of PBRM1 and/or VHL re-expression in SLR25 cells. PCNA is shown as a loading control. (C) Viability of BT549 cells following treatment with RSL3 for 16 h at the indicated concentrations, normalized to DMSO treatment group. Data are presented as mean ± SD with individual replicates shown (*n* = 3 biological replicates). Statistical significance was assessed by unpaired two-sided Student’s *t*-test comparing shPBRM1 to shScramble at each RSL3 concentration. \**P* ≤ 0.05; \*\**P* ≤ 0.01; \*\*\**P* ≤ 0.001; \*\*\*\**P* ≤ 0.0001.

**Figure S2.**
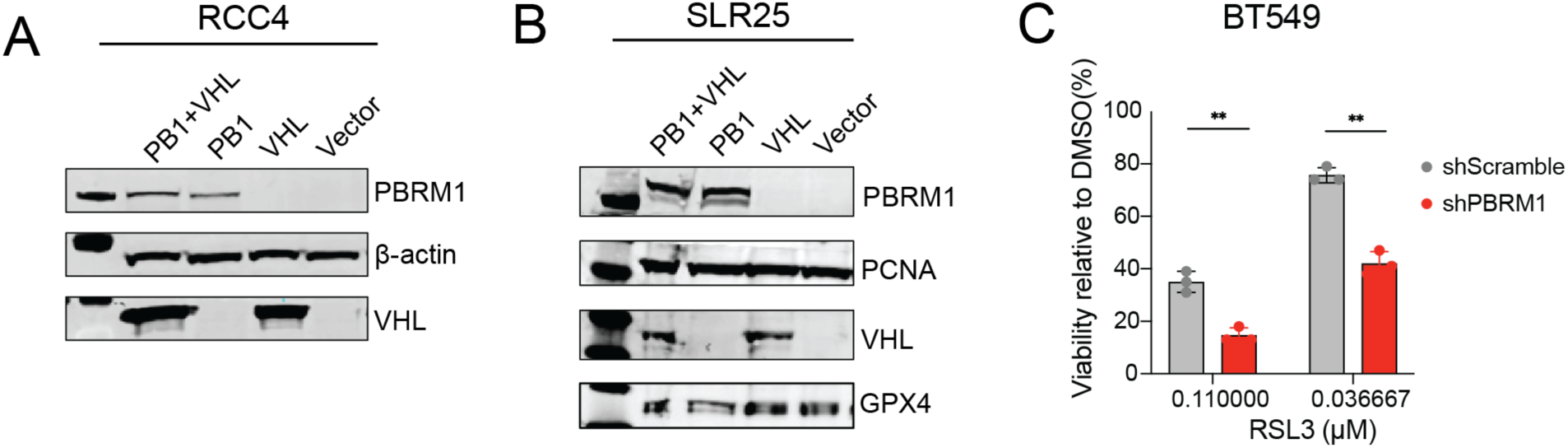
Stability of genotype effects across time and treatment and additivity of PBRM1 and VHL programs. (A-B) Volcano plots of DEGs in Caki-2 cells expressing vector (Empty), PBRM1, VHL, or PBRM1+VHL under DMSO (A) or RSL3 (0.33 μM, 10 h; T2) (B) conditions. Statistically significant DEG counts (adjusted *P* < 0.05) are indicated for each genotype. (C) Hallmark GSEA bubble plot summarizing pathway-level transcriptional differences for each reconstituted genotype relative to Empty vector under RSL3 treatment at T2. Bubble size represents −log10(adjusted *P*) and color represents NES. Only pathways with adjusted *P* < 0.05 are shown. (D) GSEA enrichment of two partial EMT gene sets — Type 2 (fibrotic/stiffness-related) and Type 3 (metastatic/migratory) — for each reconstituted genotype relative to Empty vector under DMSO conditions at T2. Bubble size represents −log10(adjusted *P*) and color represents NES. (E) Heatmap of log2 fold-changes for leading-edge genes driving Type 2 fibrotic EMT enrichment across PBRM1, VHL, and PBRM1+VHL reconstituted genotypes relative to Empty vector under DMSO conditions at T2. Color scale represents log2 fold-change relative to Empty vector. (F) Distribution of average log2 fold-changes for the full Type 2 fibrotic EMT gene set across PBRM1, VHL, and PBRM1+VHL reconstituted genotypes relative to Empty vector under DMSO conditions at T2. Vertical lines indicate the mean.

**Figure S3.**
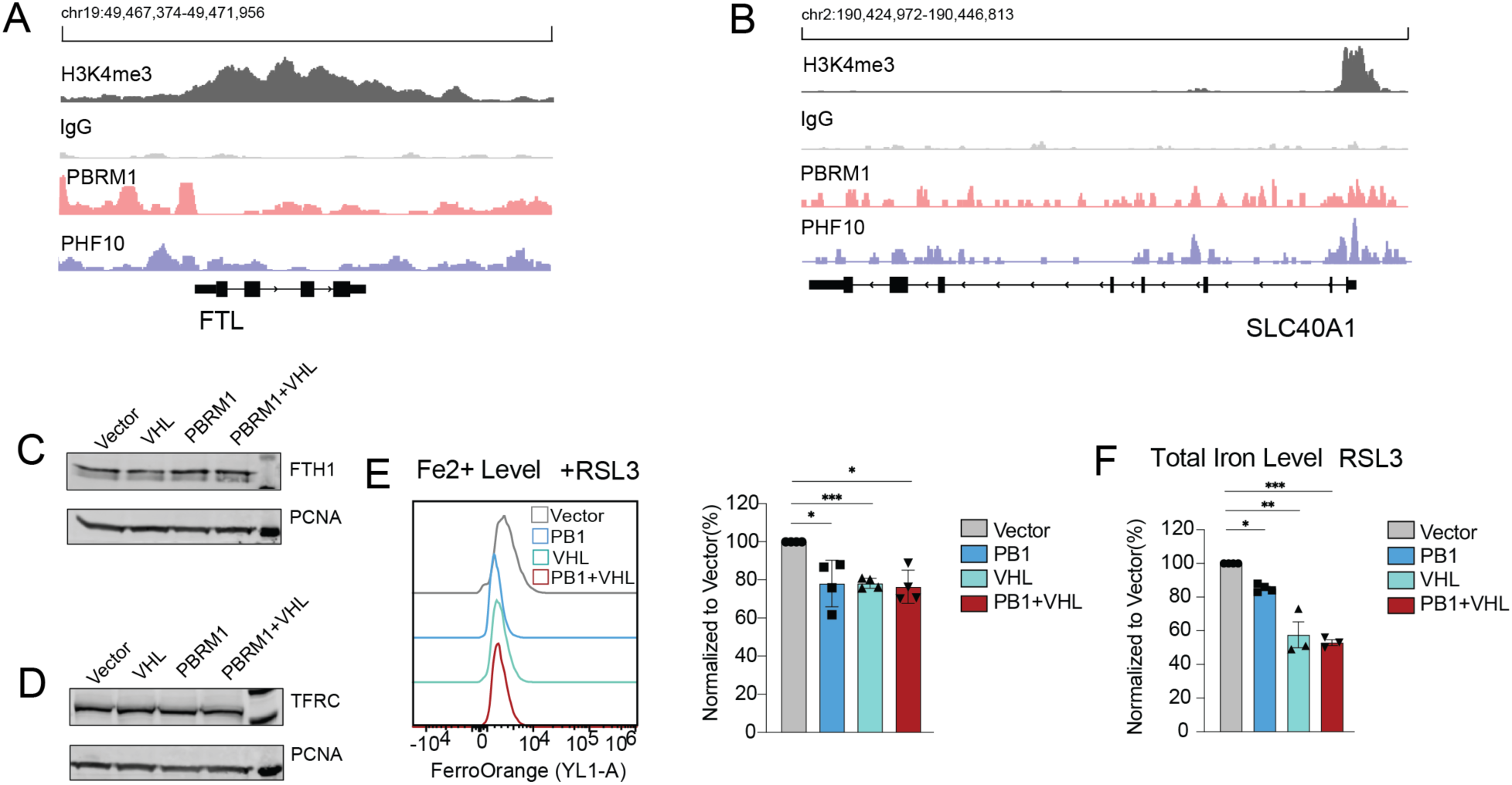
PBRM1 chromatin occupancy at iron-regulatory gene loci and corresponding protein expression. (A) CUT&RUN genome browser tracks at the *FTL* locus (chr19:49,467,374–49,471,955) in Caki-2 cells. Tracks show H3K4me3, IgG, PBRM1, and PHF10 (PBAF-specific complex subunit) occupancy. Gene structure is shown below. (B) CUT&RUN genome browser tracks at the *SLC40A1* locus (chr2:190,424,572–190,446,813) in Caki-2 cells. Tracks are displayed as in (A). (C) Immunoblot of ferritin heavy chain (FTH1) in Caki-2 cells expressing Empty vector, VHL, PBRM1, or PBRM1+VHL. PCNA serves as a loading control. (D) Immunoblot of transferrin receptor (TFRC) in Caki-2 cells expressing Empty vector, VHL, PBRM1, or PBRM1+VHL. PCNA serves as a loading control. (E) Representative flow cytometry histograms (YL1-A channel) and quantification of FerroOrange mean fluorescence intensity (MFI) in Caki-2 reconstituted cells treated with RSL3 (0.33 µM, 8 h). MFI is normalized to Empty vector. (F) Total cellular iron content (Fe²⁺ + Fe³⁺) in Caki-2 reconstituted cells under RSL3 treatment (0.33 µM, 8 h), quantified by colorimetric iron assay and normalized to Empty vector. (E-F) Data are presented as mean ± SD with individual replicates shown (*n* = 3 biologically independent replicates). Statistical significance was assessed by one-way ANOVA with Dunnett’s multiple-comparisons test using Empty vector as the reference group. \**P* ≤ 0.05; \*\**P* ≤ 0.01; \*\*\**P* ≤ 0.001; \*\*\*\**P* ≤ 0.0001.

**Figure S4.**
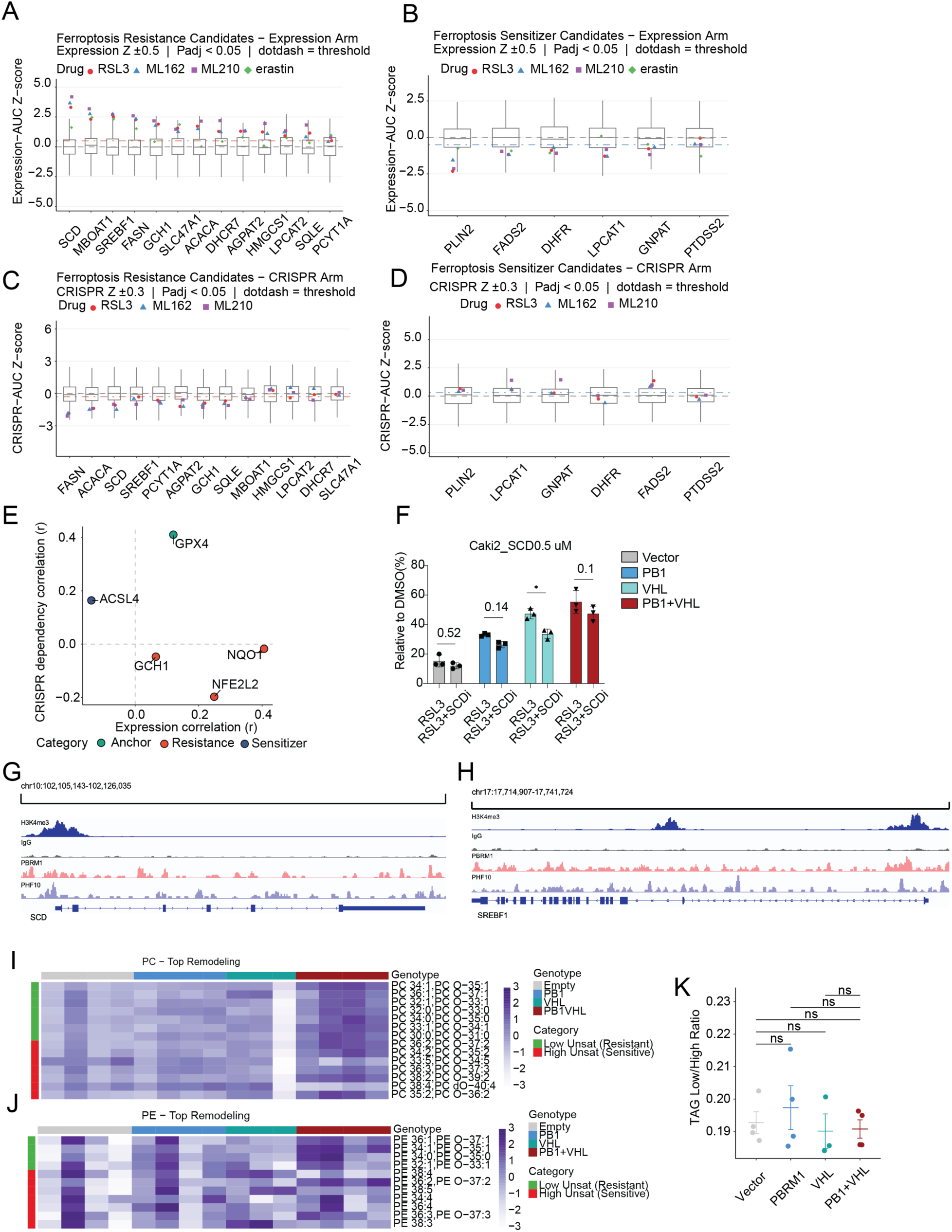
DepMap pharmacogenomic validation of candidate lipid genes and species-level lipidomic remodeling. (A) Box plots of expression-AUC Z-scores for the 13 ferroptosis resistance candidate genes across DepMap cancer cell lines. For each gene, the Pearson correlation between basal expression and ferroptosis inducer AUC was Z-scored across all compounds in the CTRP library. Individual points represent correlations for RSL3 (red circles), ML162 (blue triangles), ML210 (purple squares), and erastin (green diamonds). Dotted lines indicate the threshold of Z = ±0.5 (adjusted P < 0.05). Positive Z-scores indicate that higher gene expression correlates with ferroptosis resistance across cell lines. (B) Box plots of expression-AUC Z-scores for the six ferroptosis sensitizer candidate genes, formatted as in (A). Negative Z-scores indicate that lower gene expression correlates with ferroptosis resistance across cell lines. (C) Box plots of CRISPR-AUC Z-scores for the 13 resistance candidate genes. For each gene, Pearson correlations between CRISPR gene effect scores (Chronos) and GPX4 inhibitor AUC (RSL3, ML162, ML210) were Z-scored across all genes. Individual points represent correlations for each drug as in (A). Dotted lines indicate the threshold of Z = ±0.3 (adjusted P < 0.05). Negative Z-scores indicate that gene loss is disproportionately lethal in ferroptosis-resistant cell lines, reflecting functional dependency of the resistant state on those genes. (D) Box plots of CRISPR-AUC Z-scores for the six sensitizer candidate genes, formatted as in (B). Positive Z-scores indicate that gene loss correlates with increased ferroptosis sensitivity, consistent with a role in sustaining the ferroptosis-susceptible state. (E) Scatter plot of expression-AUC correlation (r, x-axis) versus CRISPR dependency-AUC correlation (r, y-axis) for candidate genes relative to canonical ferroptosis regulators. Points are colored by category: anchor genes (green), resistance candidates (red), and sensitizer candidates (blue). GPX4 and ACSL4 serve as positive and negative reference anchor points for expected resistance and sensitivity signals, respectively. (F) Cell viability of Caki-2 reconstituted cells following co-treatment with RSL3 (0.33 µM, 18 h) and the SCD inhibitor CAY10566 (SCDi; 0.5 µM). Viability is normalized to DMSO-treated controls within each genotype. data are mean ± SD (*n* = 3 biologically independent replicates). (G) CUT&RUN genome browser tracks at the *SCD* locus (chr10:102,105,143–102,126,035) in Caki-2 cells. Tracks show H3K4me3, IgG, PBRM1, and PHF10 (PBAF-specific complex subunit) occupancy. Gene structure is shown below. (H) CUT&RUN genome browser tracks at the *SREBF1* locus (chr17:17,714,907–17,714,724) in Caki-2 cells. Tracks are displayed as in (G). (I) Species-level heatmap of the top remodeled phosphatidylcholine (PC) lipid species across individual biological replicates for each genotype (Empty, PBRM1, VHL, PBRM1+VHL). Each row represents one lipid species annotated at sum-composition level. Rows are colored by unsaturation category: green indicates low-unsaturation species (≤1 double bond; resistance-associated) and red indicates high-unsaturation species (≥2 double bonds; sensitivity-associated). Color scale represents log2 abundance relative to Empty vector mean within each batch. (J) Species-level heatmap of the top remodeled phosphatidylethanolamine (PE) lipid species across individual biological replicates, formatted as in (G). (K) TAG low-to-high unsaturation ratio (double bonds ≤ 1 versus ≥ 2) across Empty vector, PBRM1, VHL, and PBRM1+VHL cells. Data are from lipidomics batch 2 (*n* = 4 per genotype, except VHL, *n* = 3; see Figure 7 legend and Methods). Individual replicates are shown with mean ± SEM. Statistical significance was assessed by unpaired two-sided Welch’s *t*-test relative to Empty vector. \**P* ≤ 0.05; \*\**P* ≤ 0.01; \*\*\**P* ≤ 0.001; \*\*\*\**P* ≤ 0.0001.

## Notes

### Competing Interest Statement

The authors have declared no competing interest.

